# scDiffEq: drift-diffusion modeling of single-cell dynamics with neural stochastic differential equations

**DOI:** 10.1101/2023.12.06.570508

**Authors:** Michael E. Vinyard, Anders W. Rasmussen, Ruitong Li, Allon M. Klein, Gad Getz, Luca Pinello

## Abstract

Single-cell sequencing measurements facilitate the reconstruction of dynamic biology by capturing snapshot molecular profiles of individual cells. Cell fate decisions in development and disease are orchestrated through an intricate balance of deterministic and stochastic regulatory events. Drift-diffusion equations are effective in modeling single-cell dynamics from high-dimensional single-cell measurements. While existing solutions describe the deterministic dynamics associated with the drift term of these equations at the level of cell state, diffusion is modeled as a constant across cell states. To fully understand the dynamic regulatory logic in development and disease, models explicitly attuned to the balance between deterministic and stochastic biology are required. To address these limitations, we introduce scDiffEq, a generative framework for learning neural stochastic differential equations that approximate biology’s deterministic and stochastic dynamics. Using lineage-traced single-cell data, we demonstrate that scDiffEq offers an improved reconstruction of cell trajectories and prediction of cell fate from multipotent progenitors during hematopoiesis. By imparting *in silico* perturbations to multipotent progenitor cells, we find that scDiffEq accurately recapitulates the dynamics of CRISPR-perturbed hematopoiesis. We generalize this approach beyond lineage-traced or multi-time point datasets to model the dynamics of single-cell data from a single time point. Using scDiffEq, we simulate high-resolution developmental cell trajectories, which can model their drift and diffusion, enabling us to study their time-dependent gene-level dynamics.

## Introduction

Dynamical systems underpin fundamental processes in biology and disease, including developmental differentiation and cancer. Gene expression is a standard molecular proxy to characterize cell types and states. Single-cell measurements such as single-cell RNA-sequencing (scRNA-seq) can capture snapshots of both stable cell states, as well as transient states occupied by cell subpopulations transitioning between the stable states. While individual cells are destroyed upon measurement, scRNA-seq facilitates rapid profiling of thousands of cells, enabling the development of computational strategies to infer the relationship between an observed cell state and its past and future states. These approaches facilitate the study of relationships between cell states, and between cell states and cell fates, in cell trajectories and enable new insights into the regulatory dynamics underlying developmental processes and disease.

The evolution of tools to study dynamics from single-cell molecular data has grown increasingly sophisticated, leveraging emerging techniques from machine learning and biological domain knowledge^1–6^. Trajectory inference methods have offered effective approaches for pseudotemporal ordering in low-dimensional representations of cell state, though they remain limited to correlative analyses of genes with pseudotime, restricting their ability to provide insights on the underlying mechanisms that give rise to these trajectories. RNA velocity leverages reasonable biophysical assumptions regarding nascent, mature, and degrading RNA transcripts to infer future cell states on short timescales^7,8^. Methods, including Dynamo and CellRank (and recently, CellRank2), use RNA velocity to infer long-range cell trajectories and fates^9–11^. Although these methods offer insight towards the mean drift of a cell state, they do not incorporate the possibility of cell-specific diffusion. Methods for estimation of RNA velocity from single-cell expression are also sensitive to preprocessing and smoothing operations; due to the assumptions of these operations, they may struggle to model multi-fated trajectories accurately^12^. Thus, methods that use RNA velocity as input depend on the validity of the assumptions made about transcriptional kinetics during velocity estimation.

Drift-diffusion equations have been used to model cellular dynamics from snapshot data (**Fig. 1a**). In high dimensions (>1000), such as those encountered in single-cell gene expression analyses, finding analytical solutions to partial differential equations is computationally intractable. Early frameworks to model single-cell dynamics necessitated strong assumptions about cell behavior.

**Figure 1.**
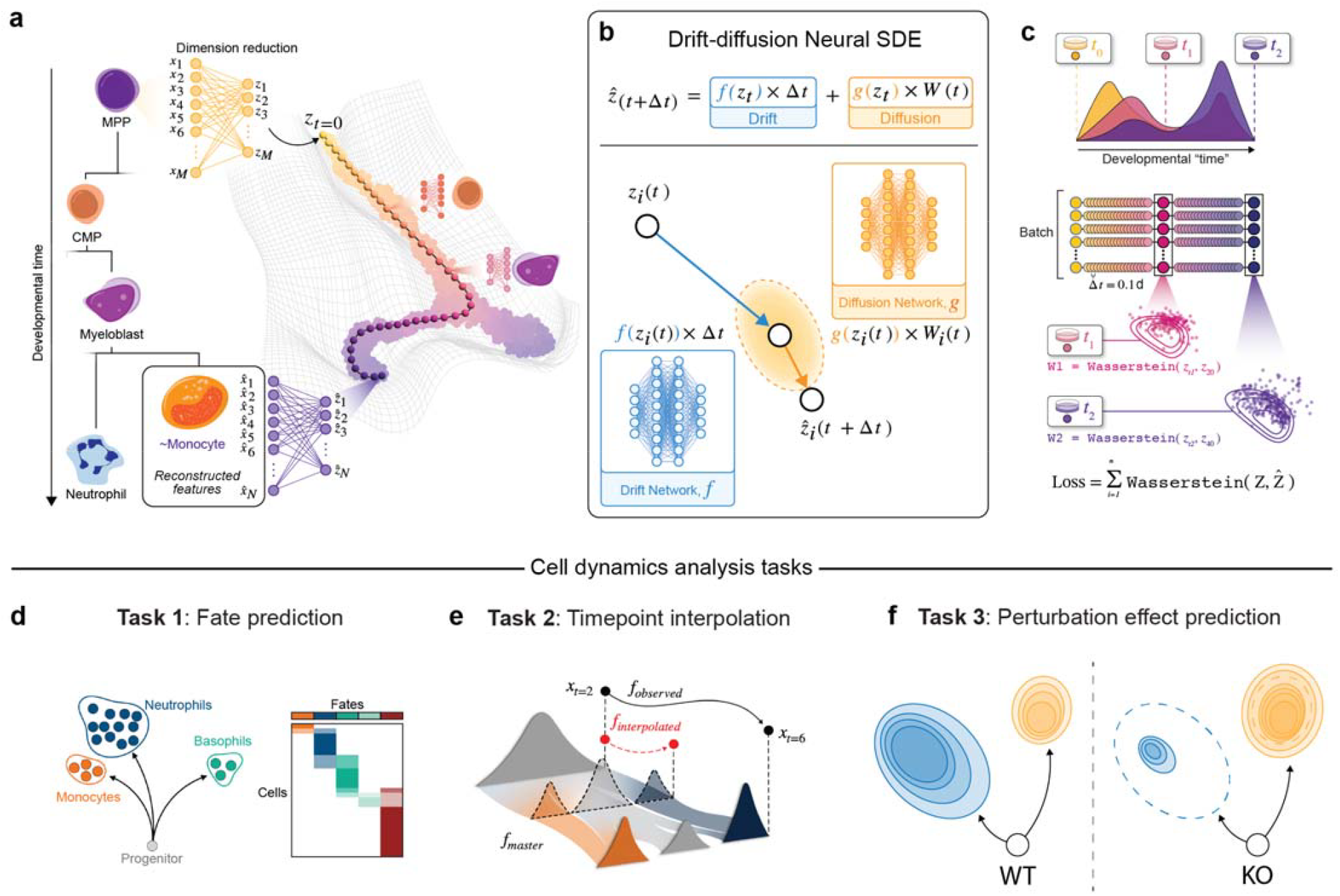
scDiffEq algorithm overview **a**. Conceptual overview of modeling a dynamical cell system such as hematopoietic development. **b**. Modeling cell drift and diffusion with neural differential equations in scDiffEq. **c**. Schematic diagram of scDiffEq training. **d**-**f**. Graphical summary of applications and analyses enabled by scDiffEq.

One of the first solutions to this framework, population balance analysis (PBA), proposed leveraging properties of spectral graph theory to model cell states at steady state via a weighted random walk through a cell neighbor graph^13^. PRESCIENT is a generative recurrent neural network that learns a drift field from coarse, time-resolved cell data using an optimal transport-based loss function. This drift field is regularized based on the assumption that cells exist in a gradient potential landscape – i.e., each forward step taken by the model is the negative gradient of the model output (potential), representing a Waddington Landscape of cell development^14,15^. Dynamo learns a smoothed vector field from noisy velocity estimates that serve as the drift term in a drift-diffusion framework^9^. Several methods, including scTour, TIGON, TrajectoryNet, and MIOFlow, have been proposed recently to use neural ordinary differential equations (neural ODEs) to describe the cell dynamics with respect to time^16–19^. WaddingtonOT, PRESCIENT, and TIGON present an improved representation of cell dynamics by explicitly modeling cell growth and death processes^6,15,17^. PBA, PRESCIENT, and Dynamo propose models using a drift-diffusion framework. However, these models restrict the diffusion to a homogeneous and isotropic Gaussian noise, treating its magnitude as a tunable hyperparameter^9,13,15^. Fixing diffusion as a dataset-level hyperparameter implies that the stochastic dynamics of individual cells are cell state-independent, thus preventing further study of the stochastic nature of gene expression as a function of cell state.

At the molecular level, stochasticity is required to facilitate the development of diverse cell types that originate from a common progenitor^20,21^. This stochasticity functions alongside more deterministic evolved regulatory mechanisms to give rise to cellular diversity observed during dynamic developmental processes. Understanding the interplay between stochastic and deterministic gene expression is essential to build interpretable models of complex cell decision-making processes. We sought to understand better how cells lean on stochasticity to make decisions. We asked where stochasticity is employed to coordinate changes to transcriptional states in time and gene expression space. Historically, differential equations have served as a workhorse of biological modeling. However, modeling complex biological systems, even in low dimensions, typically requires assumptions built on decades of empirical observations. Recent advances in deep learning, mainly neural differential equations, have provided a solution to numerically approximate dynamics governed by complex differential equations^22–24^. Neural Stochastic Differential Equations (neural SDEs) offer a direct framework to parameterize each term of a drift-diffusion equation with deep neural networks (**Fig. 1b**) ^23^. We note the independent proposal of a physics-informed neural SDE framework, PI-SDE, to model the dynamics of time-series scRNA-seq data^25^. PI-SDE was primarily demonstrated on data interpolation; here, we demonstrate the utility of using an approach that applies both data interpolation as well as fate prediction and other functional analyses, such as *in silico* perturbation, to model cell dynamics.

In this study, we build on existing models of cell dynamics, taking advantage of neural differential equations, to present scDiffEq, a deep learning framework that learns neural SDEs from embeddings of transcriptional cell states to model and study their dynamics (**Fig. 1c**). We benchmark scDiffEq against existing methods in approximating cell dynamics, using multi-time point lineage-traced scRNA-seq data (**Fig. 1d**-**f**). We note that scDiffEq predicts the dominant cell fate from multipotent progenitor cells more accurately than existing methods. We also describe scDiffEq’s ability to interpolate distributions of held-out cell populations accurately. We showcase scDiffEq’s ability to identify genes correlated with the drift and diffusion terms along simulated cell trajectories. We zoom into the granulocyte/monocyte progenitor (GMP) cell transition to neutrophils and monocytes. Along these trajectories, we recover the temporal expression dynamics of transcription factors crucial to neutrophil/monocyte fate determination. We find distinct patterns of diffusion, which we hypothesize may be necessary and sufficient for the homeostasis of the non-equilibrium transcriptional dynamics in mouse hematopoiesis. This study highlights the importance of modeling diffusion in our understanding of cellular dynamics and offers a framework for its biological interpretation.

## Results

### Learning neural differential equations with scDiffEq

scDiffEq is based on neural SDEs and is designed to accept cell transcriptional input of any dimension based on scRNA-seq data with or without time points. Contemporary methods, including PRESCIENT, use principal component analysis (PCA) as a preprocessing step to reduce the dimensionality of transcriptional cell states from thousands of measured genes to principal components^15^. For straightforward comparison, we use the first 50 principal components (PCs) of a z-scored cell-by-gene expression matrix. scDiffEq requires the annotation of an initial position from which it solves an initial value problem (IVP), to begin fitting the neural SDE describing the dynamics of the observed cell manifold. When discreetly-labeled time points are provided, scDiffEq computes the Wasserstein Distance of cells sampled from the observed cell population against those it predicts (**Fig. 1c**).

To illustrate the scDiffEq framework, in our experiments, we used a lineage-traced scRNA-seq dataset profiling mouse hematopoiesis based on the Lineage And RNA Recovery (LARRY) barcoding system, measured over three time points (days 2, 4, 6, post-barcode transduction)^26^. Despite the destructive nature of single-cell measurements, this dataset preserves state-state and state-fate relationships spanning time points. Preservation of these relationships offers an approximation of the ground truth, real-time cell dynamics. In total, the LARRY dataset comprises 130,887 scRNA-seq cell profiles: 49,302 (37.7%) of these measured cell states were successfully transduced with one of 5,864 lineage barcodes; 28,249 cells were profiled on day 2; 48,498 cells on day 4; and 54,140 on day 6. Lineage barcodes were observed in 4,638/28,249 day 2 cells (16.4%); 14,985/48,498 day 4 cells (30.9%); and 29,679/54,140 day 6 cells (54.8%). On day 2, the 4,638 barcoded cells spanned 2,672 unique barcodes. The 14,985 barcoded cells observed on day 4 spanned 4,101 unique barcodes, and the 29,679 barcoded cells observed on day 6 spanned 3,956 unique barcodes. At each time point, the most abundant barcodes occupied 8 cells, 56 cells, and 157 cells representing 0.17%, 0.38%, and 0.53% of barcoded cells, respectively.

Using this dataset, we demonstrate how scDiffEq samples cells from the day 2 population and approximates a small, discrete step “forward”, advancing at a specified interval (dt). At each annotated, observed time point (such as days 4 and 6), it simultaneously computes the Wasserstein Distance between the predicted cell population at that time point with a randomly sampled subset of the observed cells; this distance is approximated using the Sinkhorn Divergence (**Methods**). scDiffEq is then iteratively optimized to minimize the Sinkhorn Divergence between the predicted and observed cell manifolds, summed over each real time point (day 4 and day 6, using the LARRY dataset). Since scDiffEq is a generative tool that serves to create synthetic cell trajectories, these simulated cell states may appear similar to cells observed in the original scRNA-seq data, but they are unique. To facilitate several of the analyses presented in this manuscript, including fate prediction and fate perturbation screens, we map these simulated cell states to annotations of the real cells. To accomplish this, we use a nearest neighbor graph trained on the PCA space of the original data, which provides a means of mapping cell state labels (e.g., cell type annotations) to novel, simulated cells. To quantify how well the observed data might be recapitulated via simulation, we sampled an increasing number of simulated trajectories from the model and used a nearest neighbor graph to map the predicted cells to the observed cell manifold, revealing that we are able to recapitulate 74.4±1.52% of the LARRY dataset manifold from just 10,000 initial cells, and 85.5±0.99% from 50,000 initial cells (**Suppl. Fig. 1**). Once the model has converged, cell trajectories simulated in conjunction with the IVP-solver may then accurately represent a cell traversing through the learned latent time and space, enabling prediction of cell trajectories from clonal lineage families.

### Benchmarking models of single-cell dynamics with multi-timepoint, lineage-traced scRNA-seq data

Benchmark datasets paired with standardized analysis tasks are required to validate and compare predictive models. However, single-cell data generation destroys the measured cell, impeding the observation of ground truth relationships between measured cells and their true past and future states. Despite this inherently obfuscated observation of cell-cell relationships, we and others have employed the LARRY dataset as an approximation of the ground truth real-time cell dynamics (**Fig. 2a, b**). The LARRY dataset circumvents the destruction of cell trajectories by transducing multipotent progenitor (MPP) cells with lentiviral barcodes, which are heritably propagated to their daughter cells. Briefly, two days post-transduction, the MPP cells were sampled for scRNA-seq and divided into parallel wells. Each well of cells were measured via scRNA-seq at days 4 and 6 post-transduction. While not every cell sampled for scRNA-seq retained a heritable barcode, 5,864 lineage barcodes were recovered, spanning 49,302 of the 130,887 measured cells, and thus enabling the coarse reconstruction of real-time cell development in hematopoiesis over three time points (**Fig. 2c, Suppl. Fig. 2**). This dataset is among a growing collection of lineage-traced single-cell datasets that pair heritable DNA barcodes with single-cell sequencing measurements, offering temporally dependent descriptions of cell states and thereby a real-time reconstruction of the temporal dynamics including state-state and state-fate relationships ^26–33^.

**Figure 2.**
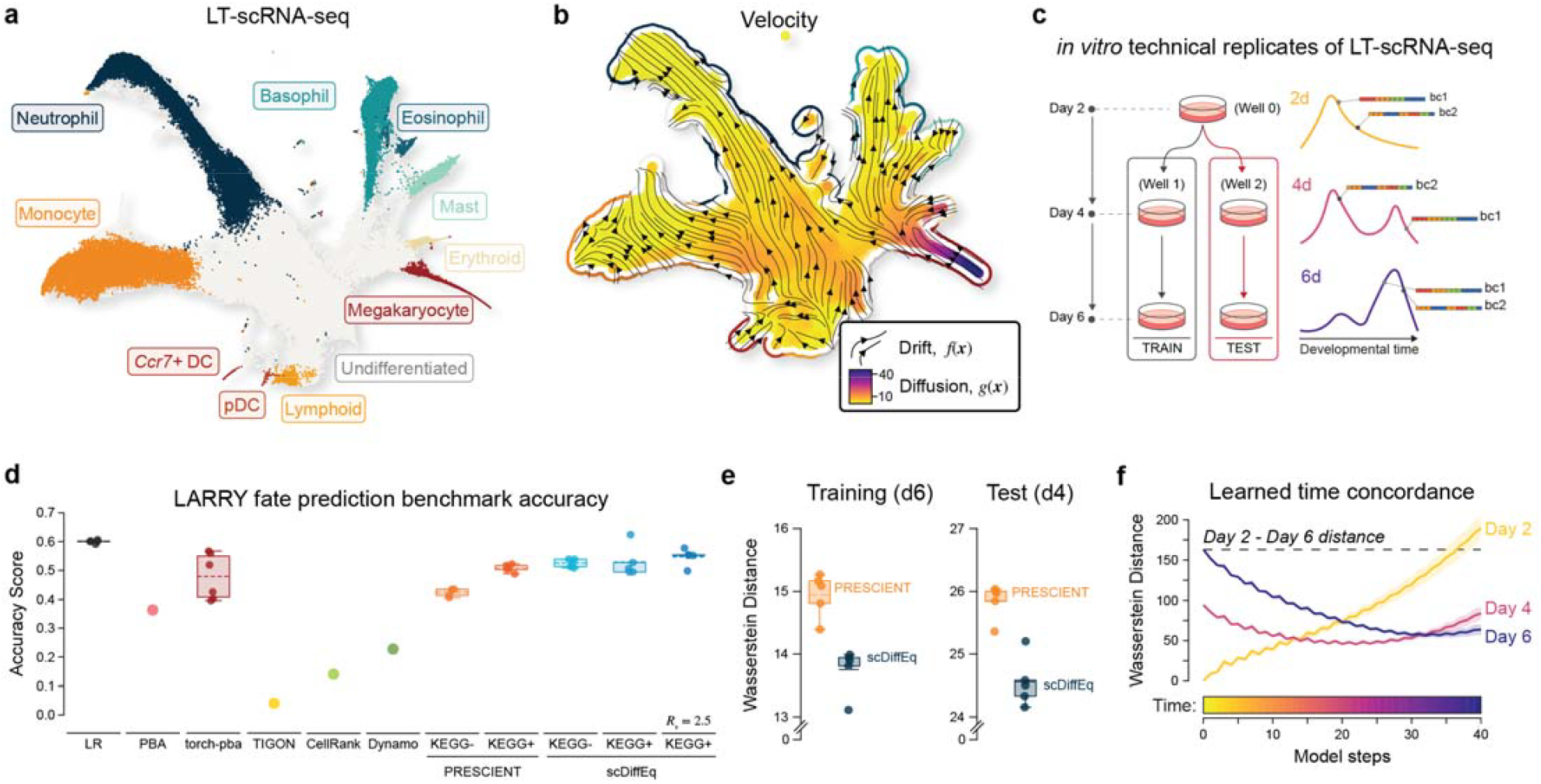
Benchmarking models of cell dynamics using the LARRY dataset. **a**. UMAP of the in vitro LARRY scRNA-seq dataset colored by cell type labels. **b**. UMAP stream plot illustrating scDiffEq-learned cell velocity decomposed into drift (vector field) and diffusion (L2Norm plotted using the colormap). **c**. Schematic overview of the lineage-tracing strategy and experimental setup of the in vitro LARRY dataset. **d**. Fate prediction accuracy for each tested method. **e**. Task 2 performance, comparing the relative ability of scDiffEq and PRESCIENT to minimize the Sinkhorn Divergence (y-axis) for the training (d6) and test (d4) sets. **f**. Sinkhorn Divergence distance (y-axis) of the predicted distribution of cells, at each discretized model time step (0.1d) against the true cell population. Horizontal dashed line indicates the distance between the observed distributions of cells at Day 2 and Day 6.

Here, we adapt and build on benchmark tasks that have previously been used in conjunction with this LARRY dataset ^9,15,17,26^ to compare the accuracy of models aimed at learning and predicting cellular dynamics on two tasks: fate bias prediction (Task 1) and prediction/interpolation of intermediate cell states at unobserved time points (Task 2). Additionally, while not benchmarked against other methods, we quantify scDiffEq’s ability to predict the relative fate consequence of perturbing progenitor cells (Task 3). For Task 1, we reasoned that for a given heritably barcoded cell observed at day 2 in the LARRY dataset, should a matching barcode be identified in another cell at one or more later time points in the dataset, we can infer that those cells are clonally related and represent an individual cell lineage. Thus, in a multipotent cell system, the barcode of a progenitor cell lineage enables a glimpse into how that lineage may or may not be biased towards formation of a specific cell fate. Taking advantage of the LARRY dataset’s ability to highlight state-fate relationships, we first benchmarked scDiffEq against several methods, including methods specialized towards modeling single-cell transcriptomic data as well as more general classification algorithms such as linear regression. The goal of the fate prediction task is to accurately infer the final “fate bias” or relative proportion of cell fates formed from a given progenitor cell, compared to the observed values tabulated for each lineage observed in the LARRY dataset (**Fig. 1d**). To prepare the LARRY dataset for each task, we followed the pre-processing procedure published in the work describing PRESCIENT (**Methods**)^15^. Briefly, for the fate prediction task (Task 1), we first segmented the dataset into train and test sets wherein all day 2 cells and the day 4 and day 6 cells from Well 1 were used as the training set. Cells in Well 2 were reserved for the test set (**Fig. 2b, Methods**). We then randomly initialized the parameters of scDiffEq using five different random seeds and trained the model as described above using the training set data. For each cell lineage, we then sampled the model, simulating 2,000 trajectories. Each trajectory began from an individual progenitor cell, generating a distribution of final states. The final states of each trajectory were annotated with a cell type label using a pre-constructed neighbor graph. These distributions of cell fates were tallied to represent the model prediction of cell fate, for each progenitor (**Methods**).

Notably, a discriminative classifier such as linear regression predicts cell fates with greater accuracy (60.0±5e-3%) than any of the single-cell focused methods that were benchmarked here (**Fig. 2d**). While a classification method offers a reasonable prediction of cell fate from the initial cell state in a 50-dimension PCA space, it does not provide information towards the underlying molecular processes and dynamics. We thus begin our benchmark with these baselines towards determining the learnable information content of scRNA-seq state descriptions that may be used to make predictions of future cell states. We also focus on the relative insights each method offers in the context of their predictive capabilities. PBA predicted cell fates with 36.3% accuracy (n=1). Torch-PBA, our PyTorch implementation of the PBA framework (**Methods**), predicted cell fates with a mean accuracy (n=5) of 53.6%. TIGON predicted cell fate with 18.0% accuracy (n=1). Dynamo, which offers in-depth analyses of generatively sampled cell trajectories, only predicted cell fates with 26.3% accuracy. CellRank, while widely-adopted, predicted cell fates with only 15.6% accuracy. Notably, Dynamo and CellRank are the only methods in this benchmark that rely on RNA-velocity estimates, using the information derived from ratios of nascent and mature mRNA transcripts. Notably, Dynamo and CellRank performed worse than all other methods featured here, except for TIGON. PRESCIENT outperformed CellRank and Dynamo, predicting cell fate with an mean (n=5) accuracy of 39.4% (without KEGG weights) and 48.7% (with KEGG weights). “KEGG weights” refer to the growth-informed weights (via KEGG gene expression signature) used in the Wasserstein Distance loss calculation as performed in previous studies using a similar optimal transport framework^6,15^. scDiffEq outperformed all methods tailored towards single-cell analysis in cell fate prediction with mean accuracies (n=5) of 52.5±1.3% (without KEGG weights) and 52.9±4.9% (with KEGG weights), representing a ∼4.2% improvement in prediction accuracy beyond the existing state-of-the-art method, PRESCIENT (**Fig. 2d**). We note that scDiffEq fate prediction performance appears to be less dependent on the use of growth weights, when compared to PRESCIENT. In general, we find that modeling non-uniform diffusion (as compared to PRESCIENT) enables us to more accurately predict fates outside of Neutrophils and Monocytes (**Suppl. Fig. 6**).

In the interest of progressing towards models with improved interpretability, we asked how the ratio of cell change (d*x*) is ascribed to the drift and diffusion fields. During model fitting, we monitor the L2Norm of both drift and diffusion for a given batch, noting a final drift:diffusion ratio of 0.37±0.14 (**Suppl. Fig. 3**). We next asked whether model regularization via an enforced target drift:diffusion ratio may provide consequential impacts to model performance or subsequent explainability. Thus, we scanned several values for such a ratio ranging from 1.0e-03 to 30.0. For each progenitor cell, we consider the entropy over 2000 simulations as a proxy of predicted trajectory multipotency (**Suppl. Fig. 5c**). Spanning the enforced ratios of drift:diffusion, we observed distinct differences in the entropy of predicted trajectories (**Suppl. Fig. 4**). While PRESCIENT effectively predicts the dominant fate for many trajectories, the corresponding cross-entropy of the fate probability vector for each cell is often significantly less accurate compared to scDiffEq (**Suppl. Fig. 7, Suppl. Fig. 8**). Importantly, revealed by comparison to the true fate bias matrix in **Suppl. Fig. 8**, we compare to several methods that do not accurately recapitulate cell fate. Using LARRY fate prediction benchmark performance as a guide, we find results to be relatively stable within drift:diffusion ratios of 1.0 to 5.0, with the best ratio observed as 2.5 (accuracy: 0.548±0.026), observing a sharp dropoff in performance at 10.0 and above (**Fig. 2d, Suppl. Fig. 5a, b**). We, therefore, enforced 2.5 as the target ratio of drift:diffusion throughout our analyses (**Methods**).

While fate prediction is informative with respect to state-fate relationships, discriminative models are not designed to recapitulate an entire cell trajectory. Both scDiffEq and PRESCIENT enable prediction of intermediate, unobserved cell states.

In Task 2, in which we benchmark interpolation of cells from a withheld time point, we compared scDiffEq to PRESCIENT, following the procedure outlined in Yeo, et. al., 2021^15^. Briefly, models are fit to the LARRY dataset using only cells from day 2 and day 6, withholding cells at day 4. Day 4 cells then serve as the test set by which models are evaluated (**Fig. 1e**). Successful reconstruction of the withheld time point was measured using the Wasserstein Distance loss function, approximated as the Sinkhorn Divergence (arbitrary units)^34^. Over five seeds, PRESCIENT was able to achieve a mean training distance of 14.94 (SEM: 0.14) and a test distance of 25.85 (SEM: 0.11). scDiffEq was able to minimize the training distance to 13.74 (SEM: 0.14) and the test distance to 24.56 (SEM: 0.16) (**Fig. 2e**). As described above, while PRESCIENT is grounded in similar assumptions to scDiffEq, it is restricted to learning functions constrained to a potential gradient and does not explicitly parameterize the diffusion term in the drift-diffusion framework. scDiffEq models, which are of comparable size to PRESCIENT in terms of the number of parameters (two fully-connected layers of 400 nodes) though unconstrained to a gradient of potential, were unable to outperform PRESCIENT. We were eventually able to improve upon the performance achieved by PRESCIENT through increased model complexity: we composed the drift and diffusion networks of the scDiffEq model for this task using two fully-connected layers of 4000 nodes and two fully-connected layers of 800 nodes, respectively.

We note a distinct advantage in the approach taken by scDiffeq and PRESCIENT to learn and subsequently simulate a latent time, from coarse, real-time measurements of cells. Using a sample model prediction from scDiffEq, we demonstrate that, over the course of a simulated population, cells beginning with zero error from their sampled d2 population move away from that population and eventually minimize their distance to the observed d4 and d6 populations at increments of 0.1d (**Fig. 2f**).

To encourage transparency and forward progress, we make our benchmark available as an open-source implementation such that the community may readily apply and evaluate new models to this benchmarking framework (**Methods**).

Model predictions are corroborated using *in silico* perturbations. The work presented in Yeo, et. al., 2021 introduced a framework for predicting the outcome of perturbing a gene in progenitor cells, given a learned model of cell dynamics^15^. Inspired by this and other frameworks such as CellOT, we next asked whether scDiffeq could simulate expected changes to biological systems under perturbed conditions^15,35^. In Task 3, we impart *in silico* perturbations over an ensemble of transcription factors (TFs) known to be involved in granulopoiesis and neutrophil development: *Lmo4, Dach1, Klf4*, and *Cebpe*. Each perturbation was introduced to the z-scored gene expression matrix of 200 randomly selected undifferentiated cells from day 2. The unperturbed, zero-centered z-score values of target genes in selected cells are then directly set to a positive value for over-expression or a negative value for perturbations that represent knockdown or knockout. For each perturbed progenitor cell, the modified gene expression z-scores were transformed into the latent space using the original PCA model - these values were used as input to scDiffEq to simulate the resulting trajectories. We compared the simulated trajectory for perturbed cells to that of an unperturbed control simulation, qualitatively noting a shift in population density from monocytes towards neutrophils, when the neutrophil TF ensemble was perturbed to z=10 (**Fig. 3a, b**). Next, we systematically profiled an array of expression perturbations spanning z-scores = 2.5, 5, 7.5, and 10 for simulated overexpression and -2.5, -5, - 7.5, and -10 for simulated gene expression knockdown. We compared the fraction of cell fates accumulated at the final time point, t=d6. In the control (z=0), the mean neutrophil fate fraction was 30.0 ±0.03% and the mean monocyte fate fraction was 27.6±0.04%. For each overexpression experiment (z = 2.5, 5, 7.5, and 10), we found a corresponding dose-dependent increase in the fraction of neutrophils predicted: 35.4±3.4%, 38.2±2.8%, 42.2±3.7%, and 48.2±3.3%, respectively (p<0.05 using Welch’s independent two-sided t-test, **Fig. 3c**). Concordantly, we observed a dose-dependent decrease in the fraction of monocytes predicted: 20.6±3.15%, 18.1±2.91%, and 13.9±2.25%, respectfully (p<0.05 using Welch’s independent two-sided t-test, **Fig. 3d**). Similarly, for simulated knockdowns, while z = -2.5 does not produce a significantly different fraction of monocytes, z = -2.5, z = -5 and z = -10 produce dose-dependent decreases to the fraction of neutrophils formed (26.6±2.7%, 25.2±2.3%, 21.2±3.4%, and 18.4±2.6%, respectively, p < 0.05). z = -5, -7.5, and -10 all produced significant corresponding increases in the fraction of monocytes formed (33.5±2.7%, 35.9±2.6%, and 38.8±2.5%, respectively, p<0.05) (**Fig. 3c, d**).

**Figure 3.**
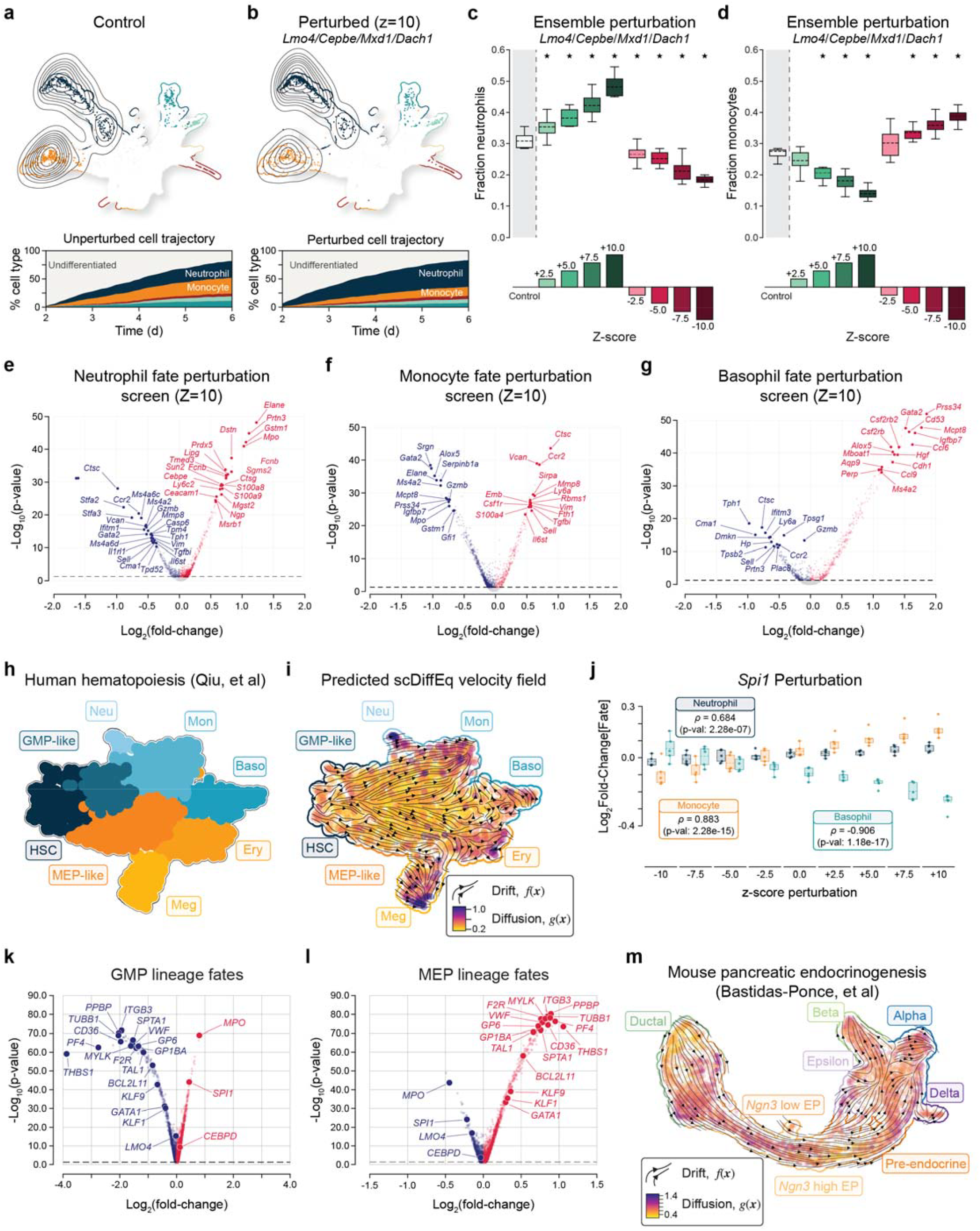
**a**. UMAP (top) highlighting the predicted distribution of cells at t = d6 under control (unperturbed) conditions and the corresponding percentages (bottom) of cell state labels at each time point in the simulated (unperturbed) trajectory. **b**. UMAP (top) highlighting the predicted distribution of cells at t = d6 under perturbation conditions and the corresponding percentages (bottom) of cell state labels at each time point in the simulated (perturbed) trajectory. **c**. Fraction of neutrophils predicted at t = 6d over varying Z-score magnitudes of the ensembled perturbation. **d**. Fraction of monocytes predicted at t = 6d over varying Z-score magnitudes of the ensembled perturbation. In both (**c**) and (**d**), Stars indicate p-value < 0.05, when compared to the unperturbed control.

There are many functional genomic screening methods to probe gene regulatory components and propose new candidate genes as regulators^36^. To expand beyond perturbing known transcription factor regulators of hematopoiesis, we next considered scDiffEq’s utility for *in silico* hypothesis generation and its potential to play a complementary role to the experimental functional genomics toolkit for studying gene regulation. Granulocyte-monocyte progenitor cells (GMPs) represent a key branch point in hematopoietic development, serving as the immediate precursor to granulocyte and agranulocyte lineages. Granulocyte lineage trajectories progress towards cell types that include neutrophils, eosinophils, and basophils, while agranulocyte lineages develop lymphocytes and monocytes. This developmental fork is explored at the gene level through an *in silico* fate perturbation screen of the LARRY dataset using scDiffEq (**Fig. 3e**-**g**).

Following the procedure described for individual genes above, for each gene in the filtered highly-variable gene set of the LARRY dataset, we impart gene expression z-score perturbations (z=10) to 200 (n=10) randomly drawn day 2 undifferentiated progenitor cells. To quantify differential fate density upon perturbation, we again employed the pre-fit neighbor graph to annotate the terminal cell states. We considered the potential for artifacts to arise from model initialization. To mitigate such variance-based artifacts and enhance generalization, we employed an ensemble approach wherein each fate perturbation screen was conducted five times, each with a distinct scDiffEq model. Each scDiffEq model was fit to the LARRY dataset from a different initialization seed. We then integrated the predicted effect sizes using meta-analysis. and combined the significance levels using Fisher’s combined probability test across the ensemble of fate perturbation screens (**Methods, Suppl. Tables 1-3**).

For each fate (monocyte, neutrophil, and basophil), we tabulated the genes that unanimously demonstrate a significant effect upon perturbation. A clustered heatmap of the effect size of these genes reveals coordinated groups of known regulatory factors that mediate the transition from GMP to granulocytes or agranulocytes (**Suppl. Fig. 10**).

For each cell fate analyzed, we highlight several known marker genes and regulatory transcription factors associated with their respective fates, or are known to govern the transition from progenitor to mature fate. For example, simulated overexpression of *Gata2* is observed to drive the largest differentially-abundant build-up of basophils (**Fig. 3g**), while simultaneously imparting the largest and most significant differential depletion of monocytes (**Fig. 3f**). Specifically directing the transition from GMP towards monocyte vs. neutrophil, we observe overexpression of *Gfi1* to be critical in enriching the accumulation of neutrophil-fated cells (**Fig. 3e**) at the cost of monocyte-fated cells (**Fig. 3f**).

Curiously, we notice the perturbation of several marker genes to seemingly confer a strong differential fate bias (**Fig. 3e-g**). While perturbation of TFs in plastic cells might be expected to impart regulatory effects, perturbation of marker genes associated with fates (e.g., *Mpo* or *Elane* in neutrophil progenitors or *Hbb* in erythroid cells) is less canonical. This disconnect may highlight a mechanistic limitation of optimal transport-based neural network models in recapitulating true causal relations. Indeed, a cell perturbed to appear more similar to a fate may effectively minimize the distance between that progenitor cell and the respective fate in PCA space, thus biasing its movement within the learned optimal transport plan.

Next, we sought to ensure that our model performs well outside of the context of the LARRY dataset. To this end, we employed the human hematopoiesis dataset described in Qiu, et al, 2022 (**Fig. 3h**)^9^. This human hematopoiesis dataset consists of 1,947 scRNA-seq profiles captured over two time points using the droplet-based metabolic labeling strategy, scNT-seq^37^. Unlike the LARRY dataset, the human hematopoiesis dataset is not barcoded for lineage tracing. We applied scDiffEq to the human hematopoiesis dataset (**Fig. 3i**), using the same hyperparameters and training specifications optimized for the LARRY dataset. We next extended the same perturbation framework used in the LARRY dataset to the human hematopoiesis dataset. Monocyte fate formation demonstrated a dose-dependent response to perturbation of *SPI1*, spanning altered perturbation levels from -10.0 to +10.0 (**Fig. 3j, Suppl. Table 4**). As in the LARRY dataset, overexpression screening (z=10) identifies several key fate-associated regulators and marker genes for the development of GMP fates and MEP fates (**Fig. 3k, l, Suppl. Tables 5, 6**). Similar to the LARRY dataset, we performed this overexpression screen over five independent replicates; for each group of fates, these replicates were strongly correlated to one another, with a minimum pairwise Pearson correlation of 0.76. (**Suppl. Fig. 11**). However, we restate the same cautionary message that it may be unrealistic to expect overexpression of markers in early progenitors to result in accumulation of the respective fates corresponding to those markers. This phenomenon is highlighted by examples including *HBB* and *HBD*. Simulated over-expression (z=+10) of these genes impart log2 fold-change effect size of 0.68 (p=6.3e-68) and 0.70 (p=6.7e-65), respectively (**Suppl. Table 5**). We expect these genes to be meaningfully expressed later in development, as markers within the MEP lineage. However, we do not realistically expect expression of these genes in HSCs to serve the stand-in role of a master regulator on which fate is dependent.

Contemporary methods that also parameterize a neural differential equation, including PRESCIENT, PI-SDE, and TIGON are restricted to using datasets consisting of multiple snapshots that span real time^15,17,25^. Thus far, we have applied scDiffEq to the LARRY dataset, which features lineage-tracing barcodes spanning three real time points as well as a human hematopoiesis dataset spanning two real time points. Here, we generalize scDiffEq beyond hematopoiesis, extending the method to a widely-used pancreatic endocrinogenesis dataset, captured at E15.5 and profiled using 3’ 10X-based scRNA-seq. (**Fig. 3m, Suppl. Fig. 12a**)^38^. Importantly, this 3,696 cell dataset is captured in a single snapshot. Since time is a required input to scDiffEq, we label each cell with a continuous developmental time (spanning 0 to 1) using CytoTRACE, a method that has been shown to faithfully recapitulate developmental time (**Suppl. Fig. 12b**)^39^. Subsequently, we relabel these values, simplifying them to 11 discrete bins, wherein the 0th bin serves as the initialization point for learning model dynamics (**Suppl. Fig. 12c**). We fit scDiffEq to the pancreatic endocrinogenesis dataset over five independent seeds, using the same hyperparameters previously used for the LARRY and the human hematopoiesis datasets (**Suppl. Fig. 12d**). Using the best checkpoint (based on validation loss) for each model, we map the learned drift-diffusion vector field to each originally observed cell in the dataset (**Fig. 3m**; **Suppl. Fig. 12e-i**). While UMAP projection-based velocity fields remain limited beyond qualitative interpretation, this dataset is widely used as an example dataset for velocity-related papers, and we observe a general relative concordance with expected results, wherein ductal cells progress towards endocrine cell fates. Here, we have demonstrated the potential for scDiffEq to serve as a critical *in silico* prediction component in functional regulation discovery toolkits, intended to complement established *in vitro* and *in vivo* approaches. Additionally, we have demonstrated an added compatibility with snapshot data, which represents most single-cell data.

### Decomposing the learned dynamics described by drift and diffusion from scDiffEq

Thus far, we have shown that scDiffEq has (i) achieved competitive performance on benchmark tasks, (ii) demonstrated its utility as a companion to model dynamics towards cell fate determination, and (iii) successfully been applied to three scRNA-seq datasets, including one without distinct time points. Next, we sought to study the trajectories that may be simulated from the scDiffEq model. We fit scDiffEq to the entire LARRY dataset over five independent seeds. We then simulated 2,000 trajectories from each of the 2,081 cell states observed in day 2 that are also heritably observed in later time points. We show three representative examples spanning the reconstruction of single-fate lineages (**Fig. 4a**), bi-fated lineages (**Fig. 4b**), and lineages with three or more fates (**Fig. 4c**). In general, this demonstration of predicting single and multiple fates from the same model represents a key limitation of other methods that focus on parameterizing a more deterministic trajectory inference model.

**Figure 4.**
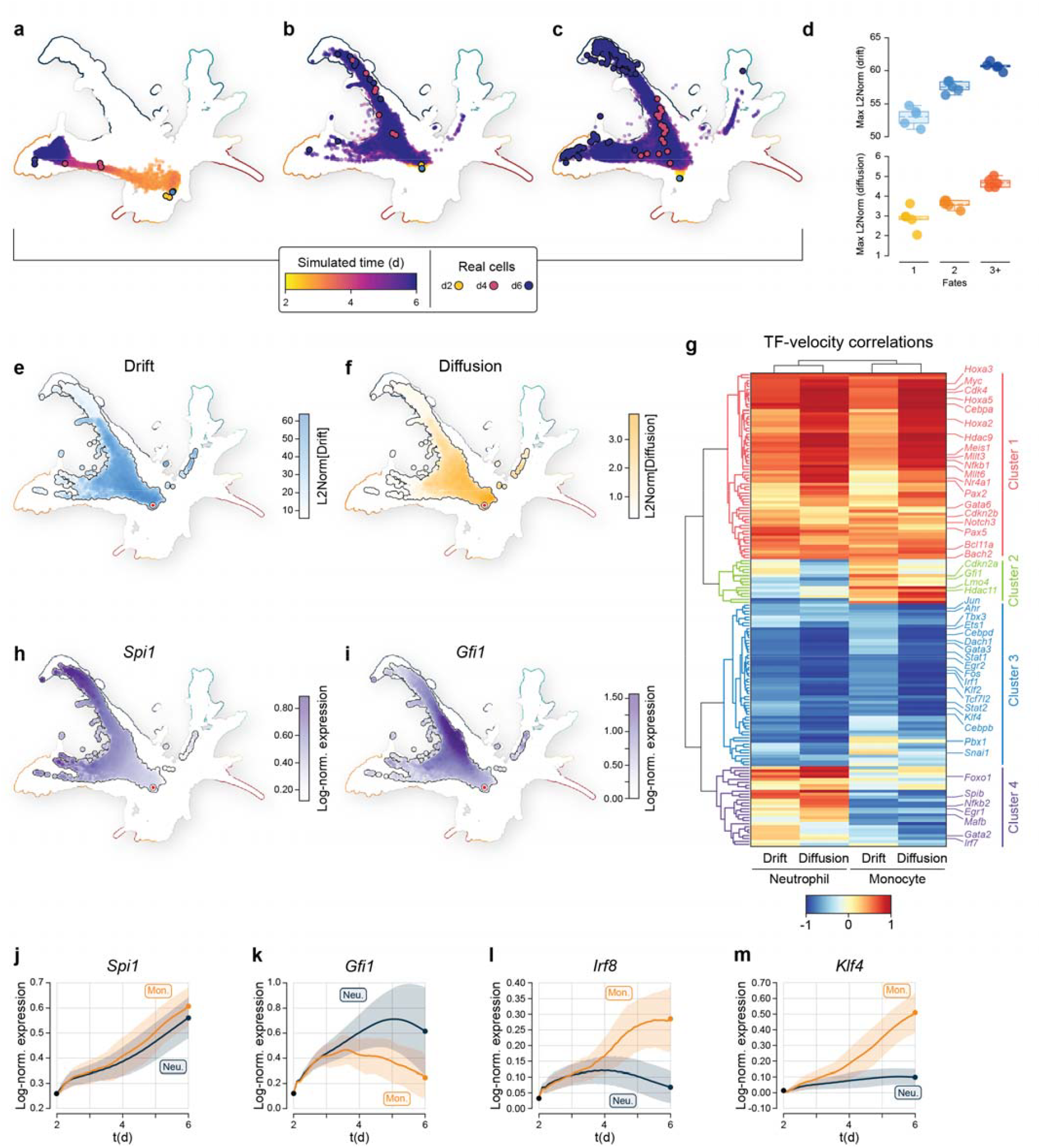
Cell-specific drift, diffusion, and time-smoothed gene expression. **a**-**c**. UMAP of an scDiffEq model simulation of a relatively mono-fated (**a**), bi-fated (**b**), and multi(3+)-fated (**c**) d2 progenitor cell from the in vitro LARRY scRNA-seq dataset. The simulation is colored according to time and plotted against the observed cell manifold. **d**. Max drift and diffusion with respect to fate multiplicity. **e, f**. UMAP projection of an scDiffEq-simulated trajectory, colored by the L2Norm of (**e**) drift and (**f**) diffusion. **h, i**. UMAP colored by the smoothed, predicted expression profile of (**h**) Spi1 and (**i**) *Gfi1*. **g**. Clustered heatmap of TF correlation to monocyte or neutrophil drift or diffusion. **j**-**m**. Simulated, fate-conditioned temporal expression profiles for (**j**) *Spi1* and (**k**) *Gfi1* (**l**) *Irf8* and (**m**) *Klf4*.

We evaluated each observed cell state of the LARRY dataset using both the independent model components: (i) the neural networks for drift and diffusion to obtain the corresponding 50-dimension drift, and (ii) diffusion vectors for each cell state. Each 50-dimension vector describes the instantaneous drift or diffusion “velocity” of each evaluated state. We summarized the magnitude of the drift and diffusion forces for each cell as a scalar value by computing the L2 norm of the 50-dimension vectors. We used a nearest neighbor graph to smooth these values and visualize using UMAP (**Suppl. Fig. 9**), noting, qualitatively, cell- and group-specific fluctuations in both drift and diffusion.

We asked whether a relationship might exist between relative cell plasticity and those cells’ drift and diffusion properties along their developmental trajectories. Briefly, we grouped lineage-traced cell trajectories categorically according to their predicted fate multiplicity: trajectories that reach a single fate; two fates; and three or more fates. We computed the L2Norm of drift and diffusion for simulated trajectories across five independent model fitting replicates. We find that the max value of the L2Norm of both the drift and diffusion along a trajectory increases monotonically as fate multiplicity categorically shifts from mono-fated trajectories (drift: 53.1±1.3, diffusion: 2.9±0.5), to bi-fated trajectories (drift: 57.6±0.8, diffusion: 3.6±0.2), to trajectories that reach three or more fates (drift: 60.7±0.6, diffusion: 4.7±0.2) (**Fig. 4d**).

From the LARRY dataset, we highlight the accurately simulated developmental trajectory of a bi-fated multipotent progenitor cell that produces both monocytes and neutrophils in roughly equal proportion, (**Fig. 4e-j**). We decompose the cell state-specific drift and diffusion along this trajectory, visualizing each trajectory using UMAP (**Fig. 4e, f**). While UMAP projections undoubtedly obscure the complexities of the real biological state space^40^, we may infer a state-dependent relationship between drift and diffusion along the neutrophil and monocyte developmental trajectories.

We isolated an example trajectory that accurately predicted (per the lineage-tracing ground truth) bi-fated neutrophil/monocyte-fated trajectories. For each trajectory, we reconstructed the gene expression by inverting the predicted PCA state vector for each simulated cell. After computing the L2Norm of the drift and diffusion for each cell in the trajectory, we compute the mean value of that drift and diffusion for each time step (dt = 0.1d), conditioned on fate. We then computed the correlation between fate-conditional drift or diffusion and reconstructed TF gene expression. Hierarchical clustering these fate-conditional correlation values uncovered four clusters (**Fig. 4g**; **Suppl. Table 7**). Cluster 1 is generally characterized by strong correlation diffusion for both monocyte- and neutrophil-fate trajectories as well as a positive, though weaker, correlation with drift for both fates. Cluster 1 contains several critical hematopoiesis genes including *Myc, Notch3, Bach2, Meis1*, and *Cebpa. Myc* is a TF and key regulator of hematopoiesis. *Cebpa* is required for the development of both monocytes and neutrophils^41^. In monocyte-fated trajectories, Cluster 2 is described by a positive correlation to drift and mixed correlations to diffusion. In neutrophil-fated trajectories, Cluster 2 hosts mixed, though mostly negative, to null correlations for both drift and diffusion. *Gfi1* and *Lmo4* are found in Cluster 2. Lmo4 maintains an interaction with Gata2. *Gata2* and *Gfi1b* are distal regulators of Myc expression in hematopoiesis^42^. *Gfi1* exhibits anti-correlation to diffusion in neutrophil-fated trajectories and a positive correlation to drift in monocyte-fated trajectories. *Gfi1* is predominantly expressed along neutrophil trajectories (**Fig. 4i, k**). *Lmo4* is associated with neutrophil-fated trajectories^15^ though strongly correlated to diffusion in monocyte-fated trajectories. Cluster 3 is predominantly characterized by negative correlations to drift and diffusion for both monocyte- and neutrophil-fated trajectories, with pronounced anti-correlation to diffusion for both fates. Genes in Cluster 3 are highlighted by several GMP development and differentiation genes, including *Cebpd, Dach1, Irf1, Klf4, Cebpb*, and *Snai1*. These correlations may be reflected through various gene expression programs. For instance, *Klf4*, while anti-correlated to both drift and diffusion in neutrophil- and monocyte-fated trajectories, exhibits greater expression in monocyte-fated trajectories, as expected, compared to neutrophil-fated trajectories (**Fig. 4m**). In contrast, also in Cluster 3, *Dach1* exhibits a similar drift-diffusion correlation profile to *Klf4*, though is more highly expressed in neutrophil-fated trajectories, compared to monocyte-fated trajectories. Cluster 4 features mixed weakly negative to strongly positive correlations to both drift and diffusion in neutrophil-fated trajectories, and predominantly negative correlations to both drift and diffusion in monocyte-fated trajectories. *Foxo1* in Cluster 4 is distinctly correlated with diffusion in neutrophil-fated trajectories, while being anti-correlated to both drift and diffusion in monocyte-fated trajectories. *Mafb and Gata2* are also found in Cluster 4.

Using scDiffEq, we model the time-dependent expression of *Spi1, Gfi1, Klf4*, and *Irf8*, separately in monocyte- and neutrophil-fated trajectories (**Fig. 4j**-**m**). *Spi1* encodes the pioneer TF protein, PU.1. *Spi1* is required for neutrophil development^41^. The GFI1 protein, encoded by *Gfi1*, is a transcriptional repressor in hematopoiesis. In early hematopoietic progenitors, *Gfi1* and *Spi1* function antagonistically through a protein–protein interaction in regulating mouse hematopoiesis^43^. Independent of fate, development beyond the GMP state is broadly dependent on *Spi1* expression^44^. Consistent with previous notions around *Gfi1* involvement in GMP development, we find sustained expression of *Gfi1* into neutrophil-annotated state space (**Fig. 4i, k**). *Spi1* expression is observed along each trajectory (**Fig. 4h, j**). Taken together, scDiffEq describes a high-resolution, dynamic portrait of *Spi1* and *Gfi1* — critical fate regulators whose antagonistic interplay shapes hematopoiesis.

## Discussion

scDiffEq is a drift-diffusion framework that leverages neural stochastic differential equations (SDEs) to capture the deterministic and stochastic dynamics of single-cell. Through rigorous benchmarking using the lineage-traced LARRY dataset, we demonstrate scDiffEq’s improved ability over contemporary methods to reconstruct cell dynamics, as measured by cell fate prediction (Task 1) and reconstruction of unseen cell populations (Task 2). We also explore scDiffEq’s ability to predict the effect *in silico* perturbations to progenitor gene expression (Task 3). We successfully applied scDiffEq across three distinct scenarios: a dataset with both lineage-tracing and multiple time points, a dataset with only two time points and no lineage tracing, and, finally, a single-snapshot dataset representative of most available scRNA-seq data. This progression demonstrates the broad applicability of our approach while establishing transparent, accessible benchmarks for future studies.

Beyond methodological improvements, scDiffEq represents a significant advance in our ability to study cell dynamics. The model’s benchmark performance suggests that explicitly modeling both deterministic (drift) and stochastic (diffusion) components better captures biological reality. By simulating cell differentiation from hematopoietic progenitors to mature granulocytes, we revealed how deterministic and stochastic changes in gene expression vary across trajectories and terminal fates. At the gene level, these simulated trajectories recapitulate experimentally observed patterns in neutrophil and monocyte lineages^26,45^. The model’s ability to distinguish between deterministic and stochastic components provides a quantitative framework for studying fate decisions, potentially resolving long-standing questions about their relative contributions in development and disease. For instance, our observation of distinct patterns in drift and diffusion along different developmental trajectories suggests that, according to the model, cells actively modulate their susceptibility to stochastic state changes during differentiation.

Several technical limitations of our approach warrant discussion. Constructing a model where the latent space is pre-computed via PCA may distort or obscure individual gene-gene relationships, preventing scDiffEq from independently discerning such relationships. This limitation is highlighted in our analysis of optimal transport (OT)-based models, where we observe markers like *Mpo* and *Elane* among the top hits in our *in silico* overexpression screen of the LARRY dataset. Similarly, in the human hematopoiesis screen, markers such as *HBD* show significant differential abundance of MEP lineage fates, though such overexpression in hematopoietic stem cells would likely not reproduce this behavior in vitro. While this behavior may be mitigated by restricting *in silico* screens to known signaling effectors, these observations underscore that tools like scDiffEq should serve a complementary role to *in vitro* and *in vivo* functional genomics approaches rather than replacing them. Additionally, while our model demonstrates scalability to large datasets (>100,000 cells), modern atlases often contain more than one million cells^46^; training neural SDEs is computationally intensive and may require additional methodological optimization.

Looking forward, we envision several directions for future development. One promising avenue is the extension of scDiffEq to integrate multiple data modalities, such as combining transcriptomic data with chromatin accessibility or protein measurements. This could provide a more comprehensive view of cellular state transitions and improve our understanding of diseases characterized by dysregulated cell state transitions, such as cancer or fibrosis. Importantly, we anticipate scDiffEq to be uniquely positioned to study the relative contributions of each modality to fate determination. Technical improvements could include alternative dimension reduction approaches that better preserve gene-gene relationships, or modifications to better regularize to the nature of stochastic decision-making. Experimental validation through targeted perturbation experiments or real-time cell tracking studies will be crucial for confirming our model’s predictions.

scDiffEq presents a flexible drift-diffusion framework for modeling the dynamics from single-cell data. We anticipate this model will extend beyond scRNA-seq to other modalities, including spatial transcriptomics or multiplexed perturbation measurements. As a generalizable method for training neural SDEs, scDiffEq establishes a foundation on which new methods for biologically-informed neural differential equation-based models may be implemented. scDiffEq is an instrumental tool for studying single-cell trajectories and cell-fate decisions and represents an important methodological step towards developing the next generation of generative models for studying biological dynamics from single-cell data.

## Methods

### Data preprocessing

Following the procedure from PRESCIENT, we filtered non-highly-variable genes, regressed out genes associated with a list of cell cycle genes^209^. For downstream analyses, we hand-selected several transcription factors and marker genes that were described in the original publication of the work or used in the analyses describing PRESCIENT^185,209^. We then scaled the log-normalized expression counts and performed PCA using sci-kit learn. As described above, we have assembled a package to reproducibly fetch, preprocess, and format the LARRY dataset and interface that dataset with models to recapitulate the benchmarking efforts we and others have used with this data. First, the data matrices and associated cell- and gene-level metadata are fetched from: https://github.com/AllonKleinLab/paper-data/tree/master/Lineage_tracing_on_transcriptional_landscapes_links_state_to_fate_during_differentiation (commit: af842ce).

### Drift-diffusion model of cell dynamics

Allow *x* ∈ *R*^*D*^ to be a vector describing the position of a cell in gene expression space. Similarly, allow *z* ∈ *R*^*D*^ to be a low-dimension representation of that cell’s position in gene expression space (e.g., 50 principal components, PCs). We assume the dynamics of a cell’s state change is represented as a drift-diffusion equation of the following form:

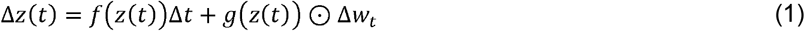

Where *f*(*z*)a n d *g*(*z*) are the drift and diffusion vector fields defin e d on *R*^*D*^, respectively. is an unobserved latent time. produces a vector describing the drift at state, which is subsequently scaled by Δ*t*. Δ*w* _*t*_ represents an increment over Brownian moti on, *W*(*t*), an i.i.d. standard spherical Gaussian. *W*_*i,j*_(*t*)∼*N*(0,1). In the case where a cell or batch of cells, is represented by a 50 PC vector, Δ*w* _*t*_ is of shape: [cells, 1], which is broadcast to [cells, 50, 1] such to facilitate element-wise multiplication against predicted the state diffusion vector, *g*(*z*(*t*)). By integrating forward in time, we may compute the future state of a cell:

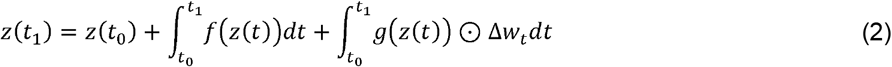

Optionally, w ithin the scDiffEq framework, one may, as demonstrate d in previous works^209^, constrain the drift field,*f*(*z*(*t*)) to the negative gradient of some potential function, *ψ* (*z*(*t*)):

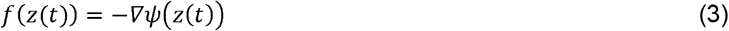

As with the drift field, *f*(*z*(*t*)), the drift field defined by is a scalar field defined on ^D^. This form of the drift-diffusion equation may be written as:

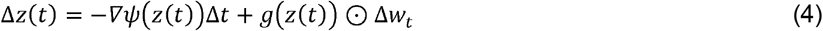

### Sinkhorn divergence

As previously implemented, we used the unbiased, entropically regularized Wasserstein distance, computed via Sinkhorn divergence as a loss metric in the reconstruction of cell populations^207,209^. Models that aim to reconstruct cell populations require a metric to compare simulated cell populations against observed cell populations. The Wasserstein distance, also known as the Earth Mover’s Distance (EMD), quantifies the minimal cost of transporting mass from one probability distribution to another. The Wasserstein distance thus provides a useful measure of the distance between two probability distributions.

Specifically, we employ the Wasserstein distance, computed via Sinkhorn divergence to quantify the distance between simulated cell populations, *υ*_*sim*_ and corresponding observed cell populations,*μ*_*obs*_. *μ* _*obs*_ and *υ*_*sim*_ are treated as probability measures on metric space, *X*. We may write the *p*-Wasserstein distance as:

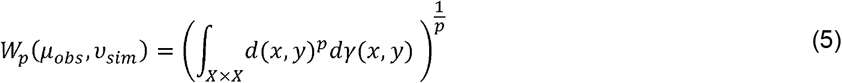

Where Π (*μ*_*obs*_, *υ*_*sim*_)denotes the set of all joint distributions, *γ* with marginals *μ*_*obs*_ and*υ*_*sim*_ ·*d*(*x,y*) is a distance metric on *X* wherein *x* and *y* represent points in *X* or the starting and ending points of mass transport in the optimal transport framework. As written, *p*-Wasserstein distance considers all possible *p*-metrics where *p* is a non-negative integer or real number that describes the type of norm used to measure the distance between points in support of their corresponding distributions. *P* = 1 describes the EMD and is more sensitive to the spread of mass distribution while *p* = 2, more frequently used in machine learning, provides a Euclidean-like measure of distance. Due to exponentiation, higher values of *p* generally lead to a distance metric more sensitive to outliers and large deviations between distributions. Here, we use a *p* =2 Wasserstein distance for its Euclidean-like distance measure properties.

As above, allow *z* ∈ *R*^*D*^ to be a vector describing a cell’s position in a low-dimension representation of gene expression space (e.g., 50 principal components). For some discrete cell population,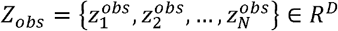 and a corresponding, model-simulated population 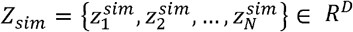, we may describe the squared Euclidean distance between vectors of *Z*_*obs*_ and *Z*_*sim*_ *as* 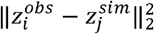.For vector *z* ∈ *R*^*D*^, this can be written as:

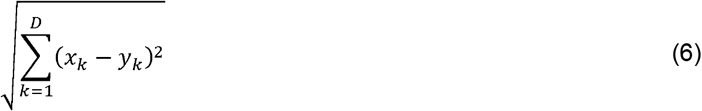

We further define the two discrete probability measures, *μ*_*obs*_ = (*Z*_*obs*_,*w*_*obs*_) ∈ *R* ^*D*^ and *υ*_*sim*_ =(*Z*_*sim*_,*w*_*sim*_) ∈ *R* ^*D*^ where *w*_*obs*_ and *w*_*sim*_ are non-negative weight vectors on the standard simplex, Δ^D^ such that observed cells weights, 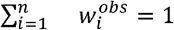 and 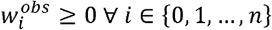. The entropically regularized Wasserstein distance between *μ*_*obs*_ and *υ*_*sim*_, with a squared Euclidean cost can be written:

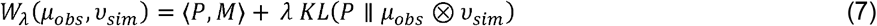

where *P* is the transport plan in the set of all transport plans *U*(*μ*_*obs*_,*υ*_*sim*_) that maps *μ*_*obs*_ to *υ*_*sim*_ and *M* is the cost matrix representing the cost of transporting mass between *μ*_*obs*_ and *υ*_*sim*_. The Frobenius inner product of the transport plan matrix *P* and the cost matrix, *M*,⟨ *P,M*⟩ describing the cost of the transport plan, is computed as the sum of element-wise products:

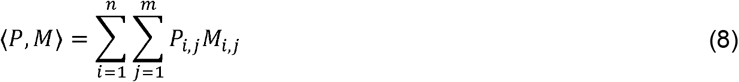

⟨ *P,M*⟩ thus represents the total cost of the transport plan matrix, *P*_*i,j*_ according to the*M*_*i,j*_ wherein we describe the cost of transporting mass from the *i*-th source to the *j*-th target. *KL* denotes the Kullback-Leibler divergence between the transport plan, *P*and *μ*_*obs*_ *⊗ υ*_*sim*_ ·*μ*_*obs*_ is an -dimension vector and *υ*_*sim*_ is a *m*-dimension vector such that *μ*_*obs*_ *⊗ υ*_*sim*_ is an *n*×*m* matrix such that the *i*-th row and *j*-th column of that matrix represent the joint probabilities *μ*_*obs*_,_*i*_*υ*_*sim*_,_*j*_, assuming the positions in *μ*_*obs*_ and *υ*_*sim*_ are independent. *λ* is a scalar regularization term applied to the *KL*-divergence, balancing the trade-off between the cost of transport and entropy in the solution.

This The marginals of the transport plan must be equal to the measures *μ*_*obs*_ and *υ*_*sim*_. Specifically, the row cost matrix, marginals of *P*_*i,j*_, the sum of each row, *i* of matrix *P*_*i,j*_ represents the total mass being transported from the *i*-th point of the observed distribution, *μ*_*obs*_ to the *j*-th point in the corresponding simulated distribution, *υ*_*sim*_. This sum must equal 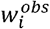 where 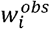 is the weight – the probability associated with the *i*-th observed point:

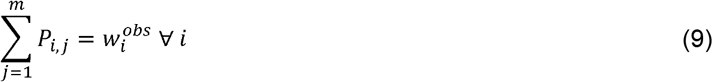

Similarly, the column marginals of *P*_*i,j*_, the sum of each column, *j* of matrix *P*_*i,j*_ represents the total mass being transported to the *j* -th point in the simulated distribution, *υ*_*sim*_. This sum must equal 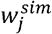 where 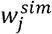 is the weight – the probability associated with the *j*-th simulated point:

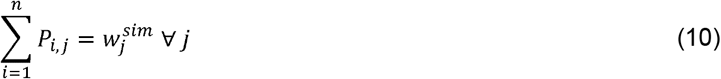

Having defined the regularized Wasserstein distance and cons trained th e marginal properties of the masses transported, the Sinkhorn divergence between two distributions *μ*_*obs*_ and *υ*_*sim*_ may be written:

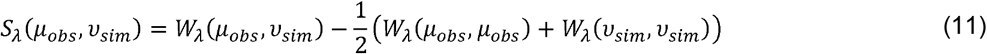

By including self-transport terms, *W*_*λ*_ and (*υ*_*sim*_, *υ*_*sim*_), the Sinkhorn divergence is equal to zero when *μ*_*obs*_ =*υ*_*sim*_. There are infinite transport plans in the set of all transport plans *U*(*μ*_*obs*_,*υ*_*sim*_). Fortunately, the solution to finding the optimal transport plan, *P*_*i,j*_ is a convex optimization problem that may be solved efficiently using a previously-implemented GPU-based solver^318^.

In scDiffEq and in similar methods, like PRESCIENT, each cell type is modeled as a particle in a distribution. The Sinkhorn divergence regularizes the optimal transport problem, enabling efficient and scalable computation of Wasserstein distances between observed and predicted distributions of cells.

### Fitting scDiffEq

In fit tin g the scDiffEq model, we aim to parameterize the terms of the above-described drift-diffusion equation, and *f*(*z*(*t*)) and *g*(*z*(*t*)) such that they accurately describe the dynamics of the observed data. Specifically, we sought to fit a NeuralSDE over the time-dependent evolution of observed cell states in *z*(*t*) from a set of progenitor cells, *z*(*t*_0_) ∈ *z*(*t*), reflective of the process shown in **Equation 2.2**. To accomplish this, we parameterized *f*(*z*(*t*)) and *g*(*z*(*t*)) as deep neural networks capable of capturing and generatively recapitulating the dynamics of cell state changes. First order discretization of **Equation 2.2** via Euler’s method (implemented by ‘torchsde.sdeint’ from Google Research) enables:

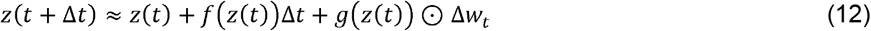

Where Δ*t* is a hyperparameter. The scDiffEq model requires the user to specify an initial *t,t*_*init*_ and a set of cell states that correspond to *z*(*t*_*init*_). Cells from *z*(*t*_*init*_) are randomly sampled in batches according to a batch size hyperparameter,. Each batch of initial cell states, *z*(*t*_*init*_) are then forward-integrated using the first-order Euler discretization shown in **Equation 2.12**, facilitated using’torchsde.sdeint’. This forward-integration produces predicted future cell states,*z*(*t* +*n*Δ*t*);*n*∈{0,1,…,*F*} where *F* Δ *t* =*T* and *T* is the final time in the dataset^267^.

If we allow *z*_*sim*_ (*t*) to represent a batch of *N* cells obtained by model simulation from *z*(*t*_*init*_) where *t* corresponds to a time annotation reflected in the observed cell dataset, we may compare the predicted states, *z*_*sim*_ (*t*) to a set of *N* cells sampled from the observed dimension-reduced gene expression states at time, *t, z*_*obs*_ (*t*).

Next, we transform *z*_*obs*_ (*t*) and *z*_*sim*_ (*t*) into probability measures,*μ*_*obs,T*_ = (*z*_*obs*_ (*t*) *w*_*obs*_ (*t*)) and *υ*_*sim*_ = (*z*_*sim*_ (*t*), *w*_*sim*_ (*t*)) where *w*_*obs*_(*t*) and *w*_*sim*_ (*t*) are growth weights derived from birth/death gene expression signatures^207,209^. We apply these gene expression signature-based growth weights only to the LARRY dataset, directly leaning on the pre-calculated values previously described^209^. Over multiple time points,*t* ∈{0,…,*T*}, the total loss used to optimize the model based on the Sinkhorn divergence of reconstructed populations and observed populations is given as:

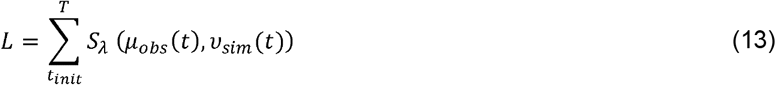

The neural networks that comprise the scDiffEq model are implemented in PyTorch. The neural network parameters are optimized through iterative forward integration, loss estimation using the above-specified Sinkhorn divergence, and backpropagation. We used the RMSProp optimizer as implemented in PyTorch. Optionally, one may also add a regularization through specifying the ratio of drift:diffusion in their respective contributions to the overall velocity. This loss is added to the *L* computed in **Equation 2.13** to yield:

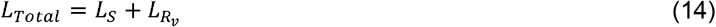

Where *L*_*s*_ represents *L* from **Equation 2.13** and 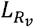 represents the regularization loss computed in **Equation 2.15** and described in the following section.

### Regularization of scDiffEq through a specified ratio of drift:diffusion

We define the ratio of velocity as:

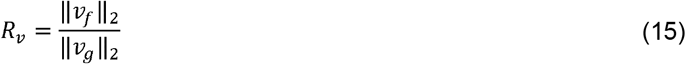

Where the numerator describes the Euclidean norm of the drift vector produced at a given cell state and the denominator describes the Euclidean norm of the corresponding diffusion vector. To impose pressure on the model to converge towards maintenance of a desired ratio of velocity, *R*_*v,target*_, we first compute *R*_*v*_ for a batch of *N* cells, *R*_*v,batch*_. We then compute the mean square error over the batch, between the target, *R*_*v,target*_ the predicted batch ratio, *R*_*v,batch*_. This error is subsequently multiplied by a constant value,*E*, which serves as a regularizing scalar by which one may differentially enforce effect of regularizing the ratio of velocity, giving:

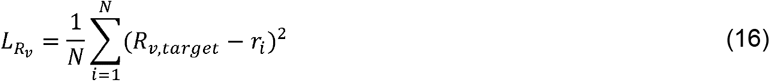

Where *r*_*i*_ is the ratio of velocity computed for a cell in *R*_*v,batch*_. Throughout this work, *E* was set to 100. As described in **Equation 2.14**, 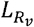 may be added to the loss derived from the Sinkhorn divergence, *L*_*s*_, to arrive at the total loss, *L*_*Total*_ over which backpropagation is performed.

### Fate prediction (Task 1) with scDiffEq

We fit scDiffEq over 250 epochs, exposing the model to cells from Well 0 (day 2) and Well 1 (day 4 and day 6), using 90% of that data for training and 10% for validation, reshuffling that split at every epoch.. Evaluation was performed by predicting the clonal fate bias of cells in Well 2 (day 4 and day 6) from the corresponding clonal progenitors in Well 0 (day 2).

### Fate prediction (Task 1) with PRESCIENT

We fit PRESCIENT, using the recommended parameters as previously described^209^. Briefly, using cells from Well 0 (day 2) and Well 1 (day 4 and day 6), PRESCIENT was trained over 2500 epochs. Evaluation was performed by predicting the clonal fate bias of cells in Well 2 (day 4 and day 6) from the corresponding clonal progenitors in Well 0 (day 2). Predicted cell states at day 6 were labeled using the pre-fit nearest neighbor classifier.

### Fate prediction (Task 1) with Population balance analysis (PBA)

To predict fate bias with PBA, we used the implementation described in its original publication. Following the procedure described in Weinreb, et al., we independently sampled 20,000 cells from the training set over five seeds^201^. PBA requires designating “source” and “sink” points in the cell manifold. Undifferentiated cells were designated as the “source”. To designate the “sink” points, we first pre-computed the potential values for each cell in the training set. For each cell fate, the cell with the minimum value of potential and its 20 nearest neighbors were designated as “sink” points. S = 10 was used as the corresponding “S” parameter for each sink point cell. Undifferentiated cells were assigned R = 0.2, while fated “sink” cells were assigned R = -0.2. Cells not designated as source or sink were assigned S = 0 and R = -1.0e-03. PBA next computes fate bias directly, producing a cell x fate bias matrix. We set the stochasticity, D = 1.0. Each cell in the training set was assigned a fate bias. To evaluate progenitor cells in the test set, we used a nearest neighbor graph built from the progenitor cells in the training set, subsequently mapping (using k = 20 neighbors) the fate biases generated for the cells in the training set to those in the test set. We also re-implemented the original PBA algorithm in PyTorch to provide a significant boost to the speed of downstream analyses (**Figure 2**).

### Fate prediction (Task 1) with Dynamo

We adapted the author-published tutorial for scRNA-seq^308^. Briefly, this approach uses PCA-transformed embeddings as input and relies on the “stochastic” model for computing expression dynamics. Importantly, the function, ’dyn.tl.gene_wise_confidence’ was used to identify and filter genes whose velocity contradicts the “direction” of cell “movement” based on user-provided initial and final states.

### Fate prediction (Task 1) with CellRank

We adapted the author-published tutorial for running CellRank using the RNA velocity kernel, which relies on pre-computing RNA velocity values using the scVelo dynamical model^307,309^. Per the tutorial, the “combined kernel” was created from a velocity kernel and connectivity kernel, weighted 0.80 and 0.20, respectively. macrostates were used to compute the absorption probabilities towards terminal states / fates.

### Fate prediction (Task 1) with TIGON

We ran TIGON using the prescribed parameters in the published protocol and a modified version of the author-provided tutorial^281^. Briefly, we fit the model using the recommended parameters described in Sha, et al. We halted the training procedure after 100 epochs. Subsequently, similar to scDiffEq and PRESCIENT, we used the model training checkpoint from epoch 100 to predict cell trajectories and fates from the 2,081 cells.

### Fate prediction (Task 1) with Linear regression

We used the scikit-learn implementation of linear regression, fitting the model to the PCA representation of the training set cells and their tabulated fate biases. We then predict the fate biases of the PCA representation of cells from the test set. We then compute accuracy scores using sci-kit learn, for all evaluated cells.

### Labeling cells using approximate nearest neighbors

Following the previously described procedure, we used a nearest neighbors classifier to label simulated cells, based on their relative position with respect to the observed manifold^209^. We used the annoy classifier from Spotify, fitting the model using all observed cells in the 50-dimension PCA space, using 10 trees, 20 neighbors, and Euclidean distance.

### Benchmark task two: recovery of a withheld time point

Following the previously described protocol, we fit both scDiffEq and PRESCIENT to the subset of data in days 2 and 6 for which lineage information was recorded^209^. scDiffEq was trained for 100 epochs. PRESCIENT models were trained for 2500 epochs. Evaluation of each model was performed by sampling 10,000 cells from day 2, with replacement, weighted by the empirically derived rate of cell proliferation and simulating one trajectory per cell. We then computed the Wasserstein distance via Sinkhorn divergence between the simulated cell populations at both day 4 and day 6 against the observed cell populations at these respective time points. PRESCIENT was evaluated at every 100 epochs. scDiffEq was evaluated at every epoch. For both models, we report the test error for the epoch with the minimum training error.

### *in silico* gene perturbations

Following the previously described protocol, we introduced perturbations to 200 cells sampled from Day 2 of the LARRY dataset^209^. For each gene perturbed, we set the z-scored expression to the indicated target value and re-transformed the scaled expression matrix using the pre-fit PCA model (to the original data). These perturbed cells were used as input to the scDiffEq model for predicting future states. We then label the model-predicted latent states using a nearest neighbor classifier fit to the original dataset. We used Welch’s independent two-sided t-test to assess significance in comparing the fractions of neutrophil and monocyte fates across conditions (perturbed vs. unperturbed). Each presented comparison was performed over 10 seeds.

## Author contributions

All authors contributed to the conceptualization of the methodology, experiments, and analyses. M.E.V. performed the M.E.V., R.L. and A.R. performed the fate-prediction benchmarking experiments. M.E.V. performed the time point interpolation experiments. M.E.V. performed the gene perturbation experiments. M.E.V. and R.L. performed the investigation of model attributes and gene-level analyses. M.E.V. wrote the software package. All authors wrote the manuscript. A.M.K. provided guidance and edited the manuscript. G.G. and L.P. provided supervision and guidance in designing the experimental strategy and funded the research.

## Acknowledgements

We graciously thank Caleb Weinreb for their assistance in using the LARRY dataset. We would like to thank Sachit Saksena, Grace Hui Ting Yeo, and David K. Gifford for their assistance in creating a benchmark comparison to PRESCIENT (Yeo et al., 2021). We would like to thank the entire Pinello and Getz Labs for thoughtful feedback and discussion throughout the preparation of this manuscript. We thank Shankara Anand for pivotal discussions that guided us towards the useful implementation of neural differential equations. We thank Arvind Ravi for useful conceptual design discussions. We thank Dongya Jia, Joseph Zhou, and Herbert Levine for useful conversations and assistance in simulating synthetic datasets for proof-of-concept experiments. We thank Mendy Miller for editorial assistance. While working on this manuscript, M.E.V. was supported by the National Institutes of Health (NIH) under the Ruth L. Kirschstein National Research Service Award (1F31CA257625) from the National Cancer Institute (NCI). L.P. was supported by the National Institutes of Health (NIH) Genomic Innovator Award (1R35HG010717-01). G.G. was partially funded by the Paul C. Zamecnik Chair in Oncology at the Mass General Cancer Center.

## Supplementary Figures

**Supplementary Figure 1.**
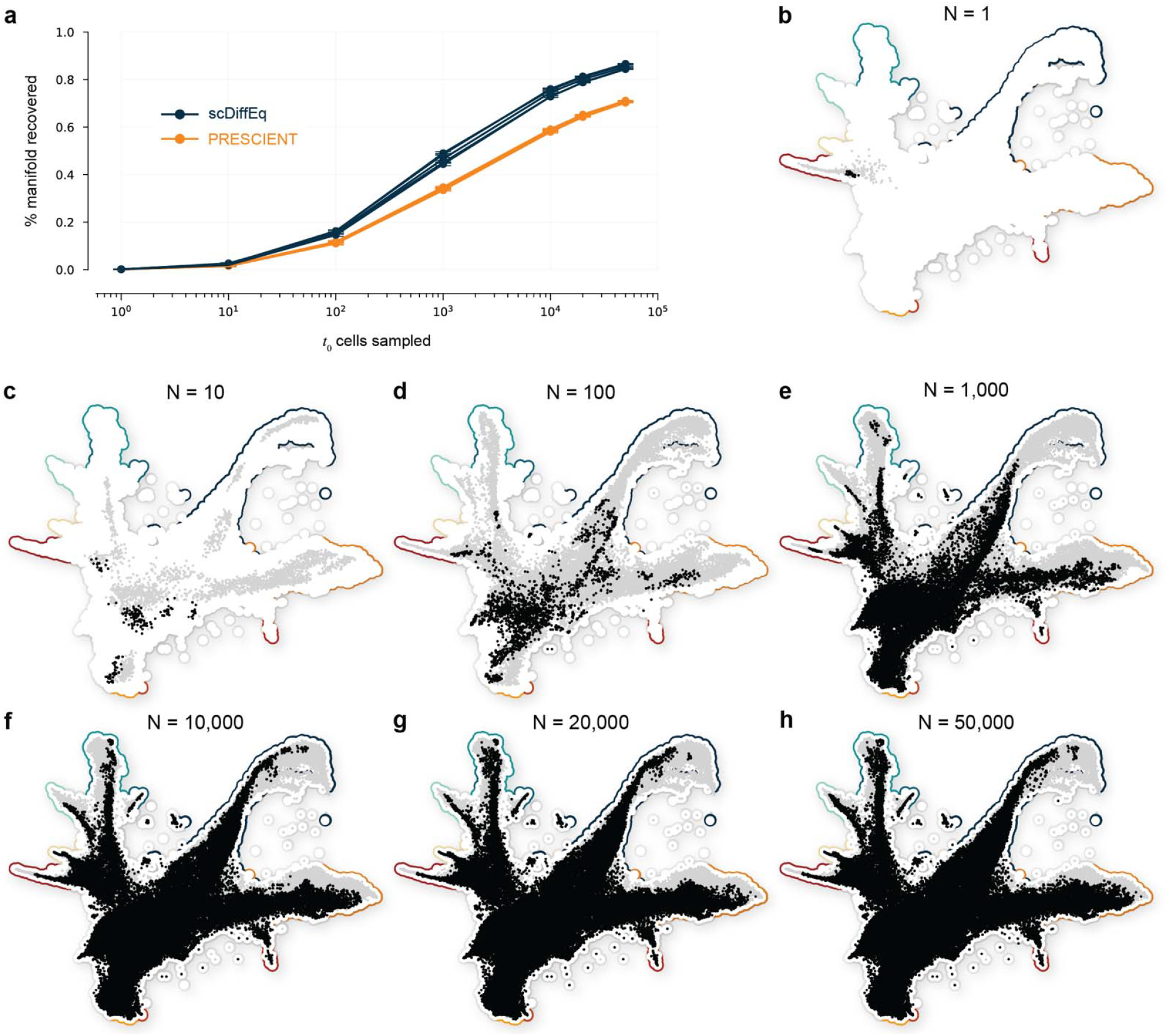
Reconstruction of the observed manifold. **a**. Percent manifold recovery, using nearest neighbor (k=20) mapping. **b**-**h**. UMAP plots showing reconstruction of the original LARRY dataset manifold (background) using model simulations (grey) from 1, 10, 100, 1000, 10,000, 20,000, and 50,000. randomly sampled initial cells, highlighted (black).

**Supplementary Figure 2.**
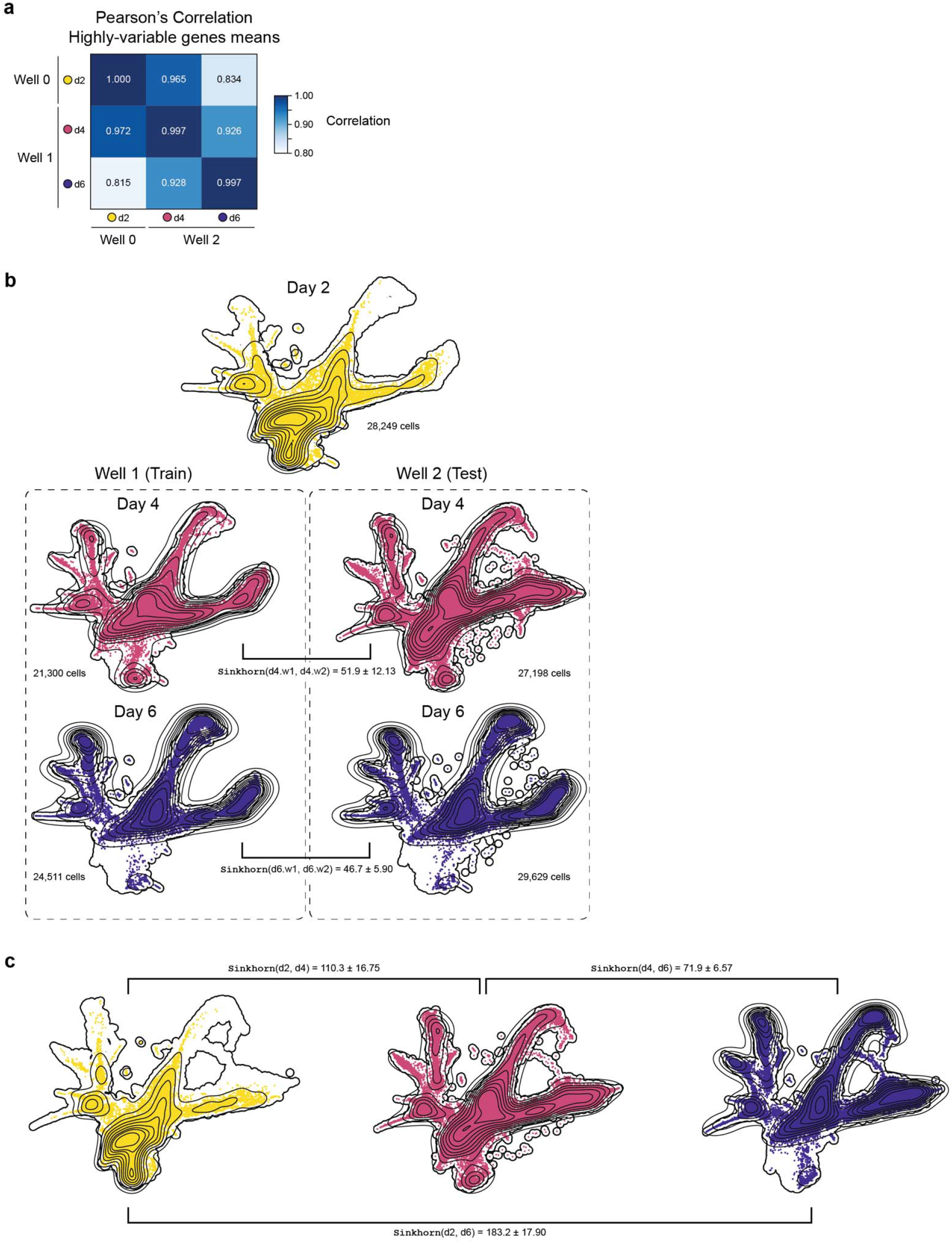
Benchmark baseline values. **a**. Pearson’s correlation coefficient of the mean of the highly-variable genes, by well. **b**. UMAP plots highlighting time-specific sub-populations of the LARRY scRNA-seq dataset, annotated with the Sinkhorn divergence sampled between wells at each time point, for fate prediction (Benchmark Task 1). **c**. UMAP plots highlighting time-specific sub-populations of the LARRY scRNA-seq dataset, annotated with the Sinkhorn divergence sampled between wells at each time point, for interpolation (Benchmark Task 2).

**Supplementary Figure 3.**
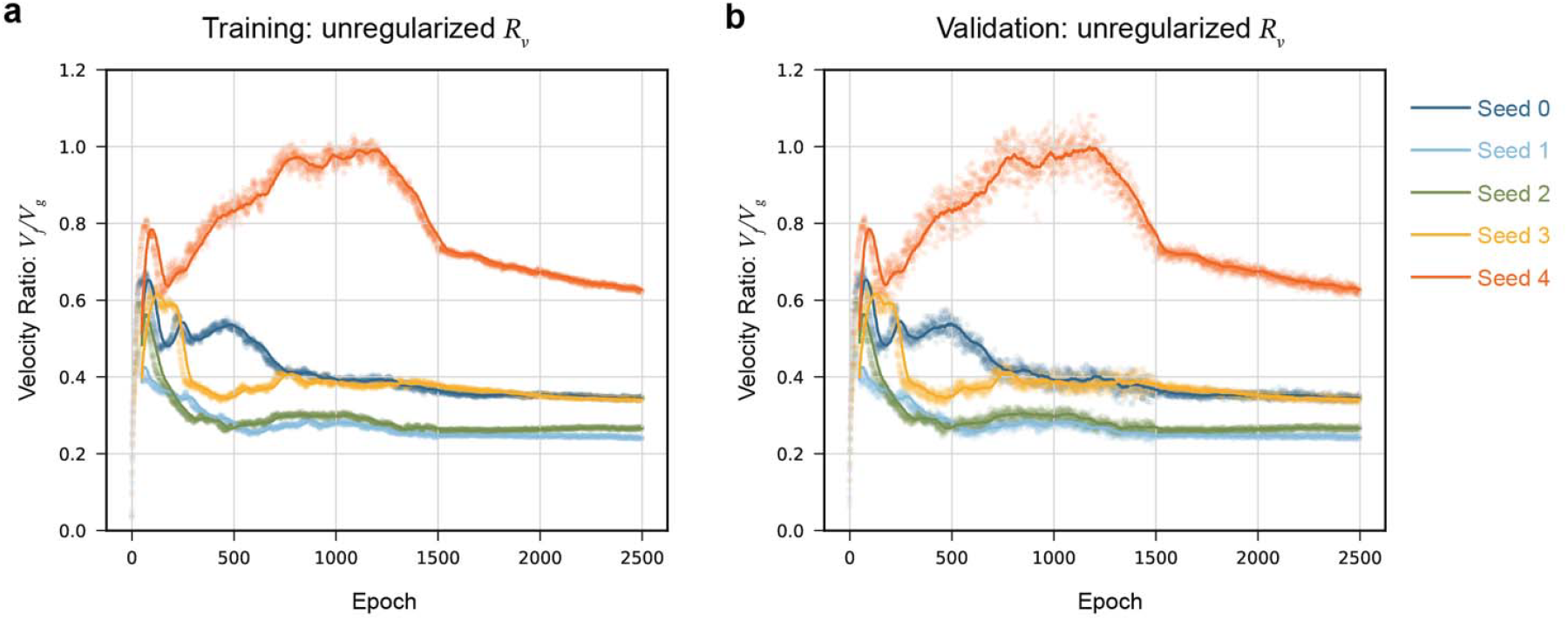
Unregularized velocity ratios observed during scDiffEq model fitting. **a**. Training **b**. Validation.

**Supplementary Figure 4.**
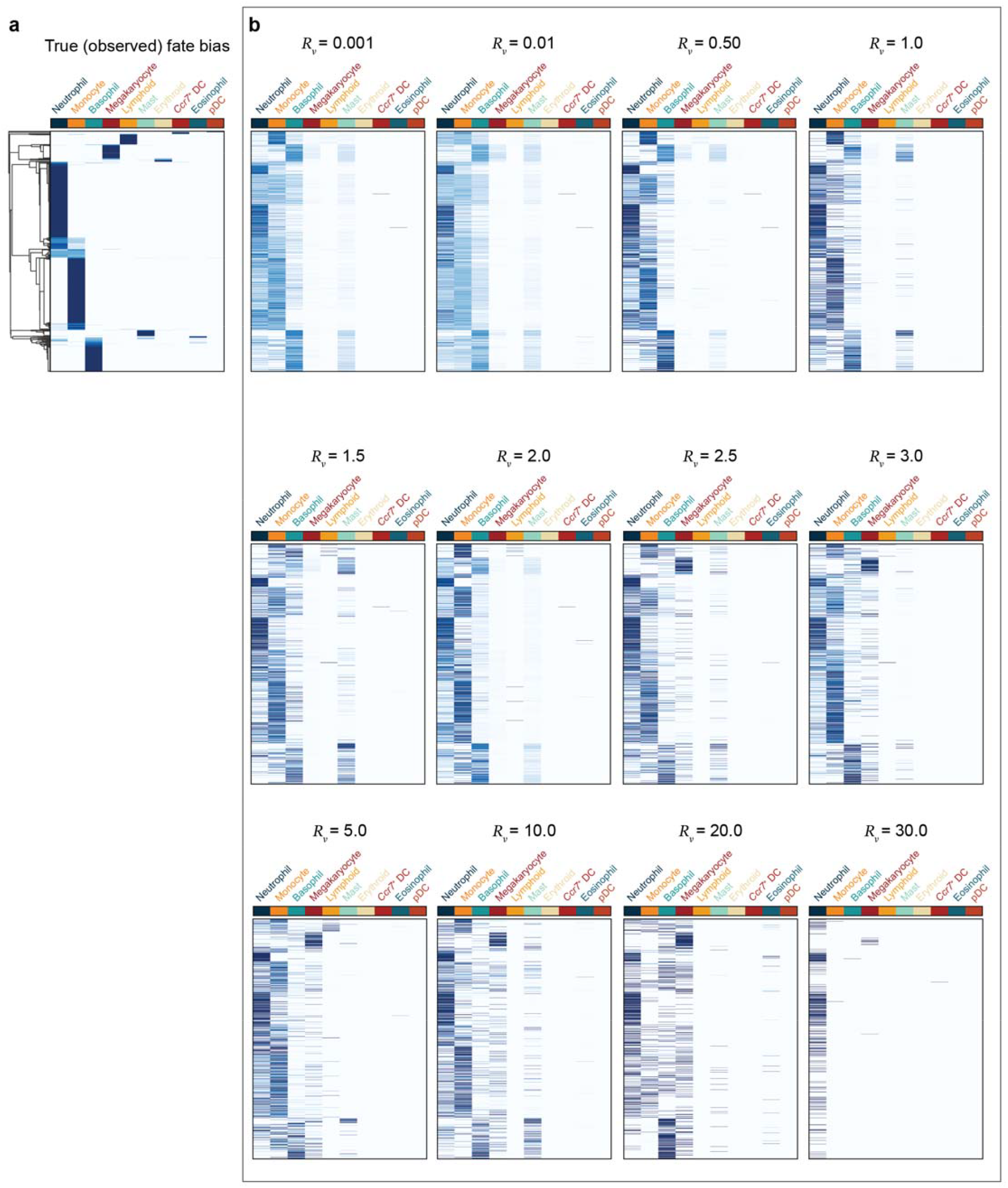
Functional consequences of enforcing a target drift:diffusion velocity ratio. **a**. Ground truth LARRY progenitor cell fate bias matrix. **b**. Predicted fate bias matrix from the best validation epoch of each imposed regularized velocity. One replicate seed shown as a representative example.

**Supplementary Figure 5.**
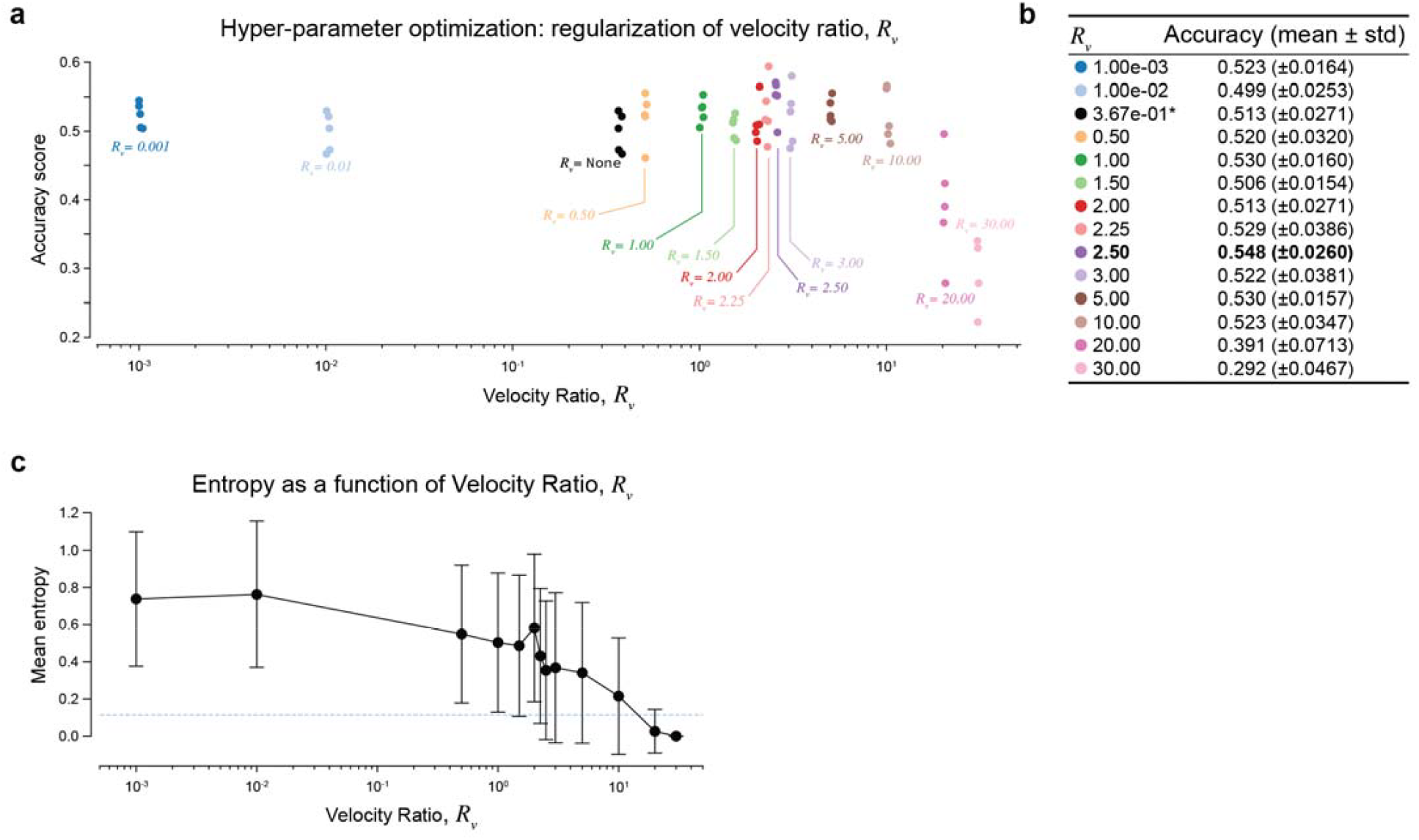
Optimization of velocity ratio. **a, b**. Hyper parameter optimization: tuning the relative ratio of enforced drift:diffusion, quantified by accuracy in prediction LARRY state-fate relationships (Benchmark Task 1). **c**. Mean per-row (progenitor cell) entropy of predicted fate as a function of the drift:diffusion velocity ratio.

**Supplementary Figure 6.**
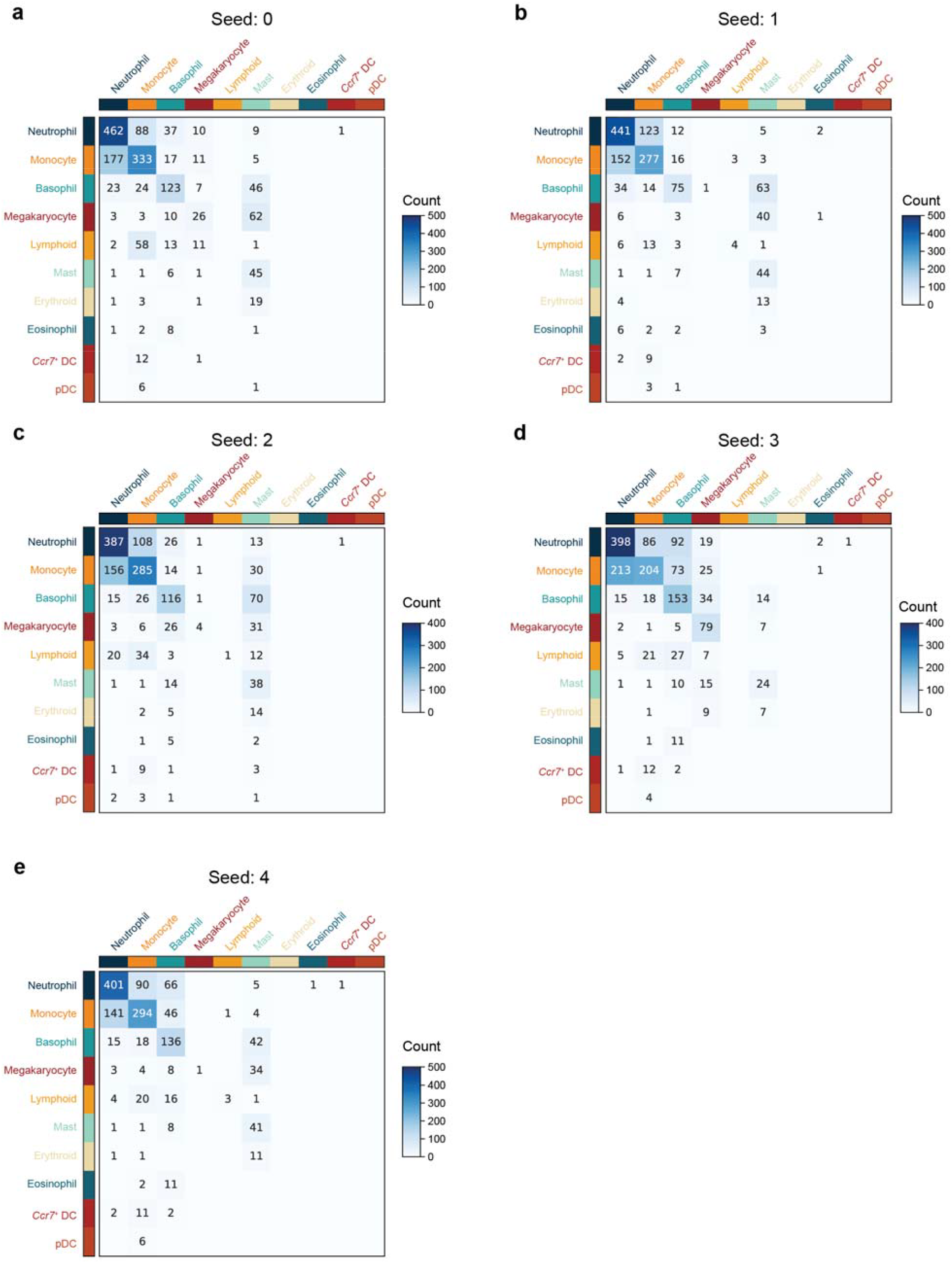
scDiffEq fate prediction confusion matrices spanning five training replicates.

**Supplementary Figure 7.**
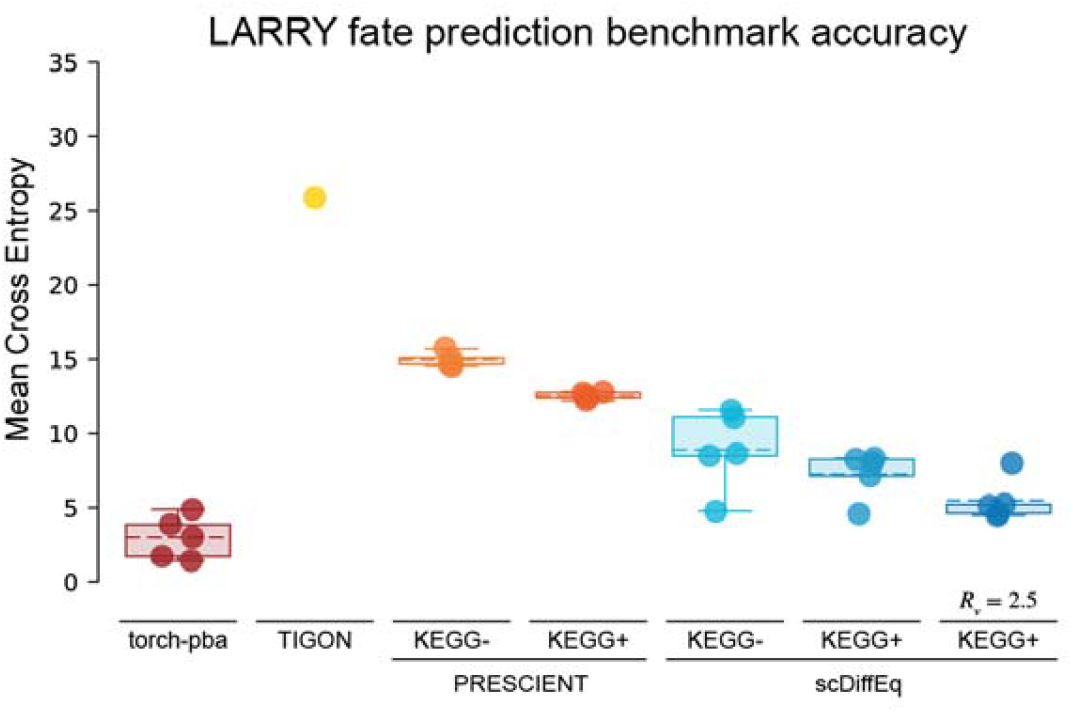
Fate prediction accuracy compared across methods by negative cross-entropy.

**Supplementary Figure 8.**
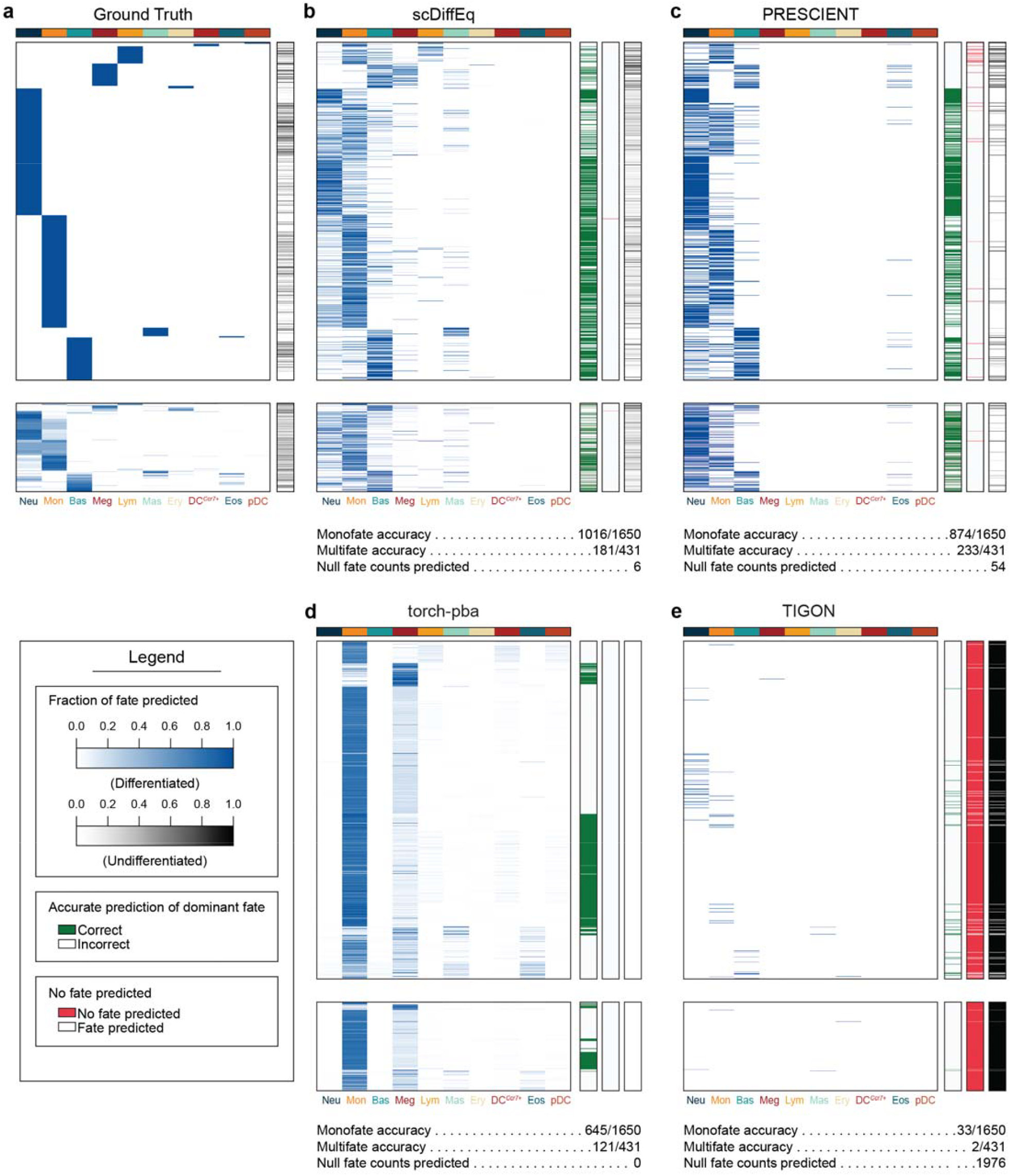
LARRY Fate bias matrices of drift-diffusion-based methods. a. Observed fate bias matrix (ground truth). Fate bias matrix for (b) scDiffEq, (c) PRESCIENT, (d) torch-pba, and (e) TIGON. For each prediction matrix, the accompanying single-column heatmaps describe whether the majority fate was accurately predicted (green), whether any fate was predicted (red for no fate predicted), and the relative proportion of undifferentiated cells predicted.

**Supplementary Figure 9.**
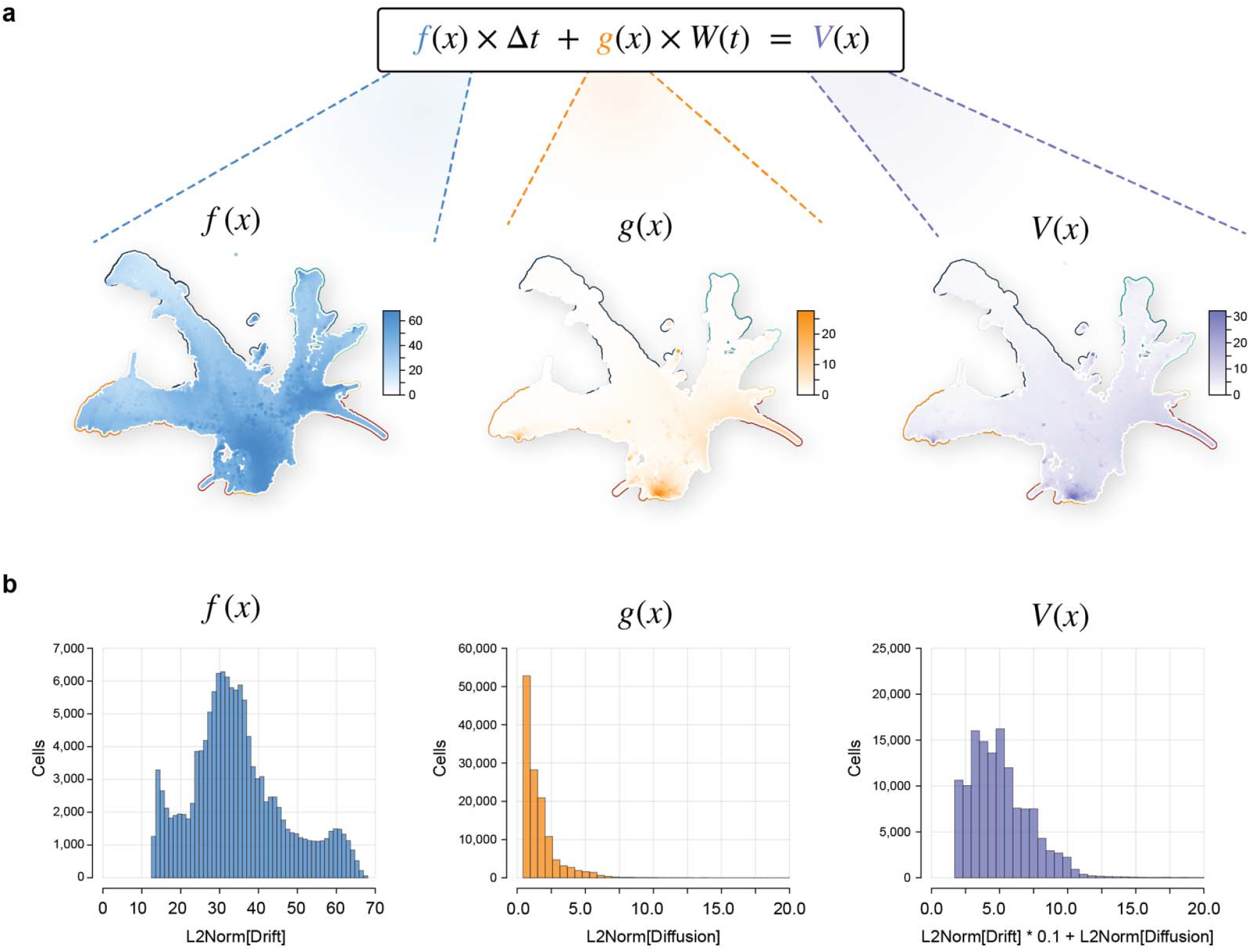
Decomposed drift-diffusion-based velocity. **a**. UMAP, colored by the decomposed drift and diffusion from a learned NeuralSDE, applied to all cells of the LARRY dataset. **b**. Histograms of drift, diffusion, and composite velocity corresponding to the cells plotted in (**a**).

**Supplementary Figure 10.**
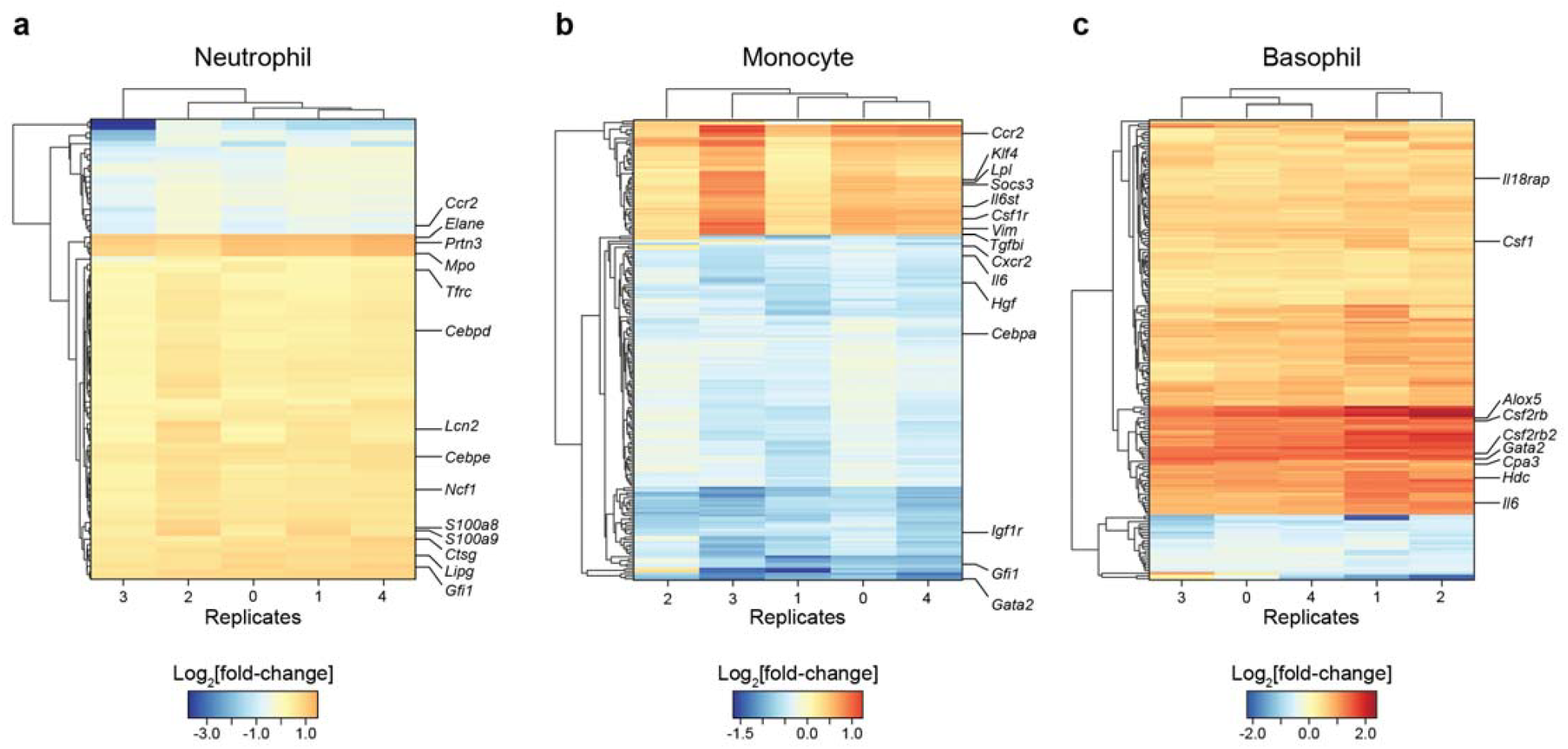
Unanimously significant genes proposed over LARRY fate perturbation screen replicates. Clustered heatmaps describing results from the fate perturbation screen of the LARRY dataset wherein each row is a gene and each column represents an independently trained model replicate and screen. Cell colors indicate the Log2Fold-change. Every gene included was below the significance threshold of p<0.5 for all five replicates. Highlighted are several genes known to be involved in the GMP lineage of hematopoiesis. A full list of the genes in each screen are listed in **Suppl. Tables 1-3**.

**Supplementary Figure 11.**
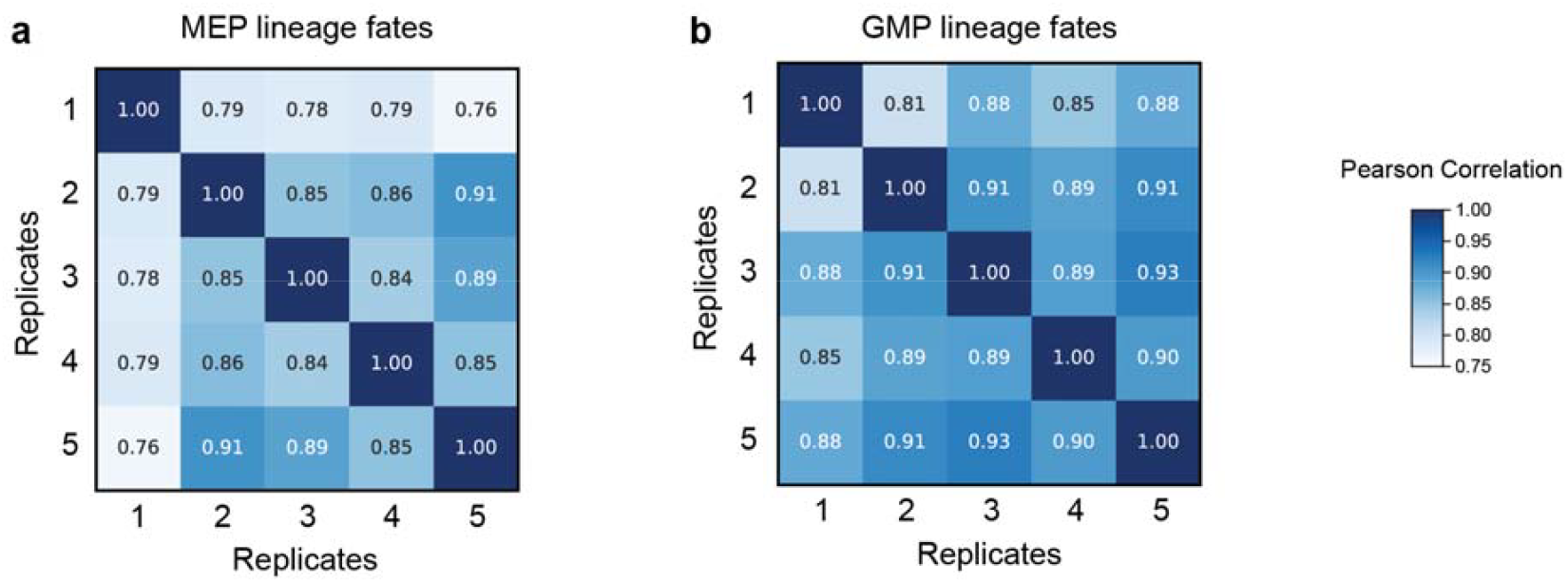
Heatmaps describing the pairwise correlation between five replicates of fate perturbation screen in the human hematopoiesis dataset. **a**. MEP lineage fate pairwise replicate correlations. **b**. GMP lineage fate pairwise replicate correlations.

**Supplementary Figure 12.**
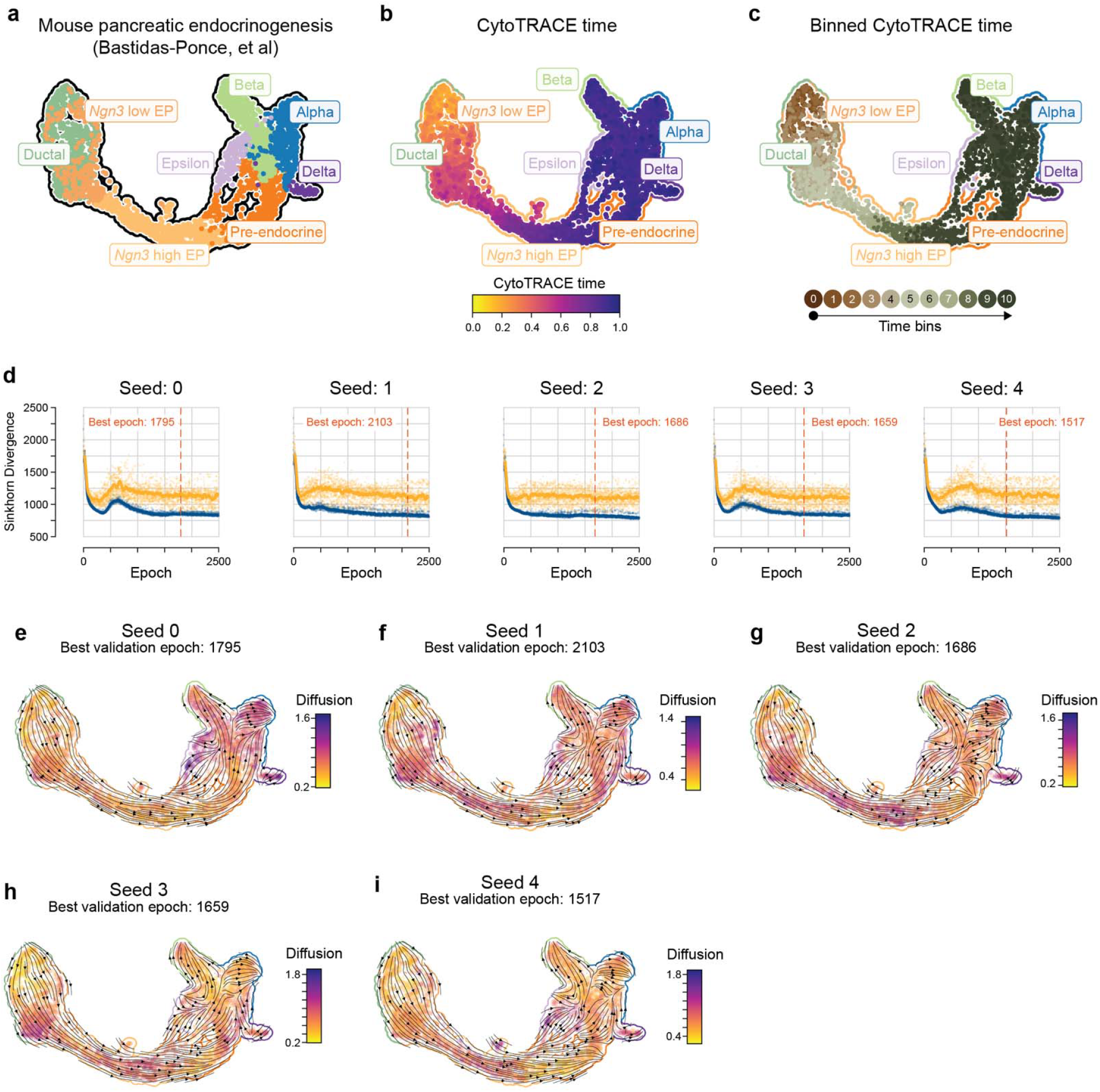
scDiffEq applied to mouse pancreatic endocrinogenesis. a-c. UMAP of mouse pancreatic endocrinogenesis, colored according to (a) cell type, (b) CytoTRACE time, (c) binned CytoTRACE time. d. scDiffEq training curves over five independent training seeds. e-i. scDiffEq velocity stream plots, using the best validation epoch from each training replicate shown in (d). Cells are colored by the L2Norm of diffusion, while the vector field represents that imparted by the learned drift field.

## References

1. Qiu, X. et al. Reversed graph embedding resolves complex single-cell trajectories. Nat. Methods 14, 979– 982 (2017).

2. Street, K. et al. Slingshot: cell lineage and pseudotime inference for single-cell transcriptomics. BMC Genomics 19, 477 (2018).

3. Chen, H. et al. Single-cell trajectories reconstruction, exploration and mapping of omics data with STREAM. Nat. Commun. 10, 1903 (2019).

4. Saelens, W., Cannoodt, R., Todorov, H. & Saeys, Y. A comparison of single-cell trajectory inference methods. Nat. Biotechnol. 37, 547–554 (2019).

5. Wang, S.-W., Herriges, M. J., Hurley, K., Kotton, D. N. & Klein, A. M. CoSpar identifies early cell fate biases from single-cell transcriptomic and lineage information. Nat. Biotechnol. 40, 1066–1074 (2022).

6. Schiebinger, G. et al. Optimal-Transport Analysis of Single-Cell Gene Expression Identifies Developmental Trajectories in Reprogramming. Cell 176, 1517 (2019).

7. La Manno, G. et al. RNA velocity of single cells. Nature 560, 494–498 (2018).

8. Bergen, V., Lange, M., Peidli, S., Wolf, F. A. & Theis, F. J. Generalizing RNA velocity to transient cell states through dynamical modeling. Nat. Biotechnol. 38, 1408–1414 (2020).

9. Qiu, X. et al. Mapping transcriptomic vector fields of single cells. Cell 185, 690–711.e45 (2022).

10. Lange, M. et al. CellRank for directed single-cell fate mapping. Nat. Methods 19, 159–170 (2022).

11. Weiler, P., Lange, M., Klein, M., Pe’er, D. & Theis, F. CellRank 2: unified fate mapping in multiview single-cell data. Nat Methods 21, 1196–1205 (2024).

12. Gorin, G., Fang, M., Chari, T. & Pachter, L. RNA velocity unraveled. PLoS Comput. Biol. 18, e1010492 (2022).

13. Weinreb, C., Wolock, S., Tusi, B. K., Socolovsky, M. & Klein, A. M. Fundamental limits on dynamic inference from single-cell snapshots. Proc. Natl. Acad. Sci. U. S. A. 115, E2467–E2476 (2018).

14. Hashimoto, T., Gifford, D. & Jaakkola, T. Learning Population-Level Diffusions with Generative RNNs. in Proceedings of The 33rd International Conference on Machine Learning (eds. Balcan, M.F. & Weinberger, K.Q.) vol. 48 2417–2426 (PMLR, New York, New York, USA, 20--22 Jun 2016).

15. Yeo, G. H. T., Saksena, S. D. & Gifford, D. K. Generative modeling of single-cell time series with PRESCIENT enables prediction of cell trajectories with interventions. Nat. Commun. 12, 3222 (2021).

16. Li, Q. scTour: a deep learning architecture for robust inference and accurate prediction of cellular dynamics. Genome Biol. 24, 149 (2023).

17. Sha, Y., Qiu, Y., Zhou, P. & Nie, Q. Reconstructing growth and dynamic trajectories from single-cell transcriptomics data. Nat Mach Intell 6, 25–39 (2024).

18. Tong, A., Huang, J., Wolf, G., van Dijk, D. & Krishnaswamy, S. TrajectoryNet: A Dynamic Optimal Transport Network for Modeling Cellular Dynamics. Proc Mach Learn Res 119, 9526–9536 (2020).

19. Huguet, G. et al. Manifold Interpolating Optimal-Transport Flows for Trajectory Inference. Adv. Neural Inf. Process. Syst. 35, 29705–29718 (2022).

20. Elowitz, M. B., Levine, A. J., Siggia, E. D. & Swain, P. S. Stochastic gene expression in a single cell. Science 297, 1183–1186 (2002).

21. Zechner, C., Nerli, E. & Norden, C. Stochasticity and determinism in cell fate decisions. Development 147, (2020).

22. Chen, R. T. Q. & Rubanova, Y. Neural ordinary differential equations. Adv. Neural Inf. Process. Syst. (2018).

23. Kidger, P., Foster, J., Li, X. & Lyons, T. J. Neural SDEs as Infinite-Dimensional GANs. in Proceedings of the 38th International Conference on Machine Learning (eds. Meila, M. & Zhang, T.) vol. 139 5453–5463 (PMLR, 18--24 Jul 2021).

24. Kidger, P. On Neural Differential Equations. arXiv [cs.LG] (2022).

25. Jiang, Q. & Wan, L. A physics-informed neural SDE network for learning cellular dynamics from time-series scRNA-seq data. Bioinformatics 40, ii120–ii127 (2024).

26. Weinreb, C., Rodriguez-Fraticelli, A., Camargo, F. D. & Klein, A. M. Lineage tracing on transcriptional landscapes links state to fate during differentiation. Science 367, (2020).

27. VanHorn, S. & Morris, S. A. Next-Generation Lineage Tracing and Fate Mapping to Interrogate Development. Dev. Cell 56, 7–21 (2021).

28. Alemany, A., Florescu, M., Baron, C. S., Peterson-Maduro, J. & van Oudenaarden, A. Whole-organism clone tracing using single-cell sequencing. Nature 556, 108–112 (2018).

29. Biddy, B. A. et al. Single-cell mapping of lineage and identity in direct reprogramming. Nature 564, 219– 224 (2018).

30. Bowling, S. et al. An Engineered CRISPR-Cas9 Mouse Line for Simultaneous Readout of Lineage Histories and Gene Expression Profiles in Single Cells. Cell 181, 1693–1694 (2020).

31. Frieda, K. L. et al. Synthetic recording and in situ readout of lineage information in single cells. Nature 541, 107–111 (2017).

32. Raj, B. et al. Simultaneous single-cell profiling of lineages and cell types in the vertebrate brain. Nat. Biotechnol. 36, 442–450 (2018).

33. Spanjaard, B. et al. Simultaneous lineage tracing and cell-type identification using CRISPR-Cas9-induced genetic scars. Nat. Biotechnol. 36, 469–473 (2018).

34. Feydy, J. et al. Interpolating between Optimal Transport and MMD using Sinkhorn Divergences. in Proceedings of the Twenty-Second International Conference on Artificial Intelligence and Statistics (eds. Chaudhuri, K. & Sugiyama, M.) vol. 89 2681–2690 (PMLR, 16--18 Apr 2019).

35. Bunne, C. et al. Learning single-cell perturbation responses using neural optimal transport. Nat. Methods 20, 1759–1768 (2023).

36. Przybyla, L. & Gilbert, L. A. A new era in functional genomics screens. Nat. Rev. Genet. 23, 89–103 (2022).

37. Qiu, Q. et al. Massively parallel and time-resolved RNA sequencing in single cells with scNT-seq. Nat. Methods 17, 991–1001 (2020).

38. Bastidas-Ponce, A. et al. Comprehensive single cell mRNA profiling reveals a detailed roadmap for pancreatic endocrinogenesis. Development 146, (2019).

39. Gulati, G. S. et al. Single-cell transcriptional diversity is a hallmark of developmental potential. Science 367, 405–411 (2020).

40. Chari, T. & Pachter, L. The specious art of single-cell genomics. PLoS Comput. Biol. 19, e1011288 (2023).

41. Giladi, A. et al. Single-cell characterization of haematopoietic progenitors and their trajectories in homeostasis and perturbed haematopoiesis. Nat Cell Biol 20, 836–846 (2018).

42. Liggett, L. A. & Sankaran, V. G. Unraveling Hematopoiesis through the Lens of Genomics. Cell 182, 1384–1400 (2020).

43. Dahl, R., Iyer, S. R., Owens, K. S., Cuylear, D. D. & Simon, M. C. The transcriptional repressor GFI-1 antagonizes PU.1 activity through protein-protein interaction. J. Biol. Chem. 282, 6473–6483 (2007).

44. Orkin, S. H. & Zon, L. I. Hematopoiesis: an evolving paradigm for stem cell biology. Cell 132, 631–644 (2008).

45. Rodriguez-Fraticelli, A. E. et al. Single-cell lineage tracing unveils a role for TCF15 in haematopoiesis. Nature 583, 585–589 (2020).

46. Regev, A. et al. The Human Cell Atlas. Elife 6, (2017).

